# Structural basis for Vipp1 membrane binding: From loose coats and carpets to ring and rod assemblies

**DOI:** 10.1101/2024.07.08.602470

**Authors:** Benedikt Junglas, David Kartte, Mirka Kutzner, Nadja Hellmann, Ilona Ritter, Dirk Schneider, Carsten Sachse

**Affiliations:** Ernst-Ruska Centre for Microscopy and Spectroscopy with Electrons, ER-C-3/Structural Biology, Forschungszentrum Jülich, 52425 Jülich, Germany; Department of Biology, Heinrich Heine University, Universitätsstr. 1, 40225 Düsseldorf, Germany; Department of Chemistry, Biochemistry, Johannes Gutenberg University Mainz, Germany; Institute of Molecular Physiology, Johannes Gutenberg University Mainz, Germany

**Keywords:** IM30, Vipp1, ESCRT-III, helical reconstruction, cryo-EM, membrane remodeling, membrane tubulation, carpet

## Abstract

Vipp1 (also known as IM30) is essential in most oxygenic photoautotrophic organisms. It is involved in membrane remodeling and fusion and is critical for thylakoid membrane biogenesis and maintenance. Vipp1 has recently been identified as a member of the ESCRT-III superfamily of membrane remodeling proteins, albeit it still is elusive how Vipp1 interacts with and finally remodels membranes. Here we present a series of cryo-EM structures of cyanobacterial Vipp1 interacting with bacterial membranes: first, we solved seven structures between 5 and 7 Å resolution of three unique helical and four types of stacked-ring assemblies engulfing membranes, and, second, using sub-tomogram averaging, we determined three ∼20 Å resolution structures compatible with previously observed carpet structures at three different membrane curvatures. By analyzing ten additional unique structures of N-terminally truncated Vipp1, we could show that helix α0 is essential for membrane tubulation and forms the membrane anchoring domain of Vipp1. Using a conformation-restrained Vipp1 mutant, we were able to reduce the structural plasticity of Vipp1 assemblies in the presence of lipids and determined two structures of Vipp1 at 3.0 Å resolution, resolving the molecular details of membrane anchoring and intersubunit contacts of helix α0. Our data reveal the molecular details of how Vipp1 interacts with membranes, showing membrane curvature-dependent structural transitions from carpets to rings and rods, some of which are capable of inducing and/or stabilizing high local membrane curvature triggering membrane fusion.

**Summary:** Bacterial ESCRT-III family member Vipp1 forms membrane-bound coats, carpets, ring complexes, stacked-ring assemblies and helical tubes capable of internalizing lipids and inducing high membrane curvature.

## Introduction

Members of the phage shock protein A (PspA) family have recently been identified as bacterial ESCRT-III proteins by structural and phylogenetic analyses (Gupta et al., 2021; Junglas et al., 2021; Liu et al., 2021). Similar to their eukaryotic counterparts (*i.e.* for *Saccharomyces cerevisiae* Vps2, Vps20, Vps24, Snf7, Vps60, Did2, and Ist1), PspA family proteins are known to form large homooligomeric complexes, such as rings or helical rods, and their proposed physiological functions include membrane maintenance and remodeling. A prominent member of the PspA family is the vesicle-inducing protein in plastids 1 (Vipp1) that is also known as IM30 (inner membrane-associated protein of 30 kDa). Vipp1 is conserved and essential in most oxygenic photoautotrophic organisms and is thought to have been transferred from cyanobacteria to the chloroplasts of algae and plants (Carlton and Baum, 2023; Kroll et al., 2001; Vothknecht et al., 2012; Westphal et al., 2001). Vipp1 binds to negatively charged membranes and membrane binding depends on stored curvature elastic stress (Heidrich et al., 2016; McDonald et al., 2015). Membrane binding has been suggested to be mainly mediated by Vipp1’s N-terminus, although N-terminally truncated Vipp1 constructs have been shown to be capable of membrane binding (McDonald et al., 2017, 2015; Thurotte and Schneider, 2019). Vipp1 appears to have two functions: first in membrane maintenance, *i.e.,* during heat or light stress, and second in thylakoid membrane (TM) biogenesis and/or remodeling (Gutu et al., 2018; Hennig et al., 2015; Junglas and Schneider, 2018; Siebenaller et al., 2019; Zhang et al., 2016, 2012, 2014). The precise structural mechanism underlying both functions, as well as Vipp1’s membrane-bound structure, are still enigmatic.

The electron cryo-microscopy (cryo-EM) structures of *Synchecystis* sp. and *Nostoc punctiforme* Vipp1 ring complexes in the absence of membranes revealed that the Vipp1 monomer structure contains six α-helices connected by short loops (Gupta et al., 2021; Liu et al., 2021; Schlösser et al., 2023), while a predicted 7^th^ C-terminal α-helix, discriminating Vipp1 from PspA (Hennig et al., 2017; Vothknecht et al., 2012), could not be resolved. Ring complex structures with variable diameters and rotational symmetry have been determined in the absence of membranes (*Synechocystis*: C14-C18 (Gupta et al., 2021); *Nostoc*: C11-C17 (Liu et al., 2021)). The rings assemblies consist of six to seven stacked layers that are formed from radial-symmetrically arranged monomers. The conformations of the monomers change from layer to layer, leading to a tapered dome-like shape of the ring complexes (Gupta et al., 2021; Liu et al., 2021). The monomers differ at the hinge regions between helices α3 and α4 (hinge 2), α4 and α5 (hinge 3), and in the length of α4. In the ring complexes, a non-canonical nucleotide binding site has been identified in the layers with the smallest diameters of each ring (Gupta et al., 2021). Although multiple studies confirmed the NTPase activity of Vipp1, it does not seem to critically affect the formation of ring complexes, membrane binding, and/or membrane fusion (Junglas et al., 2020b; Ohnishi et al., 2018, 2022; Siebenaller et al., 2019).

Apart from ring complexes, Vipp1 forms polymeric assemblies, such as elongated stacked-ring assemblies, rods/tubes of helical organization, and two-dimensional carpets, although none of these structures have been solved in molecular detail (Junglas et al., 2020a; Saur et al., 2017; Theis et al., 2019). Interestingly, Vipp1 helical assemblies have been shown to engulf lipid membranes *in vitro* and have also been found to interact with the TM *in vivo* (Gupta et al., 2021; Theis et al., 2019). Vipp1 carpets were observed to form on solid-supported bilayers *in vitro* and possibly also *in vivo* during high-light stress (Gutu et al., 2018; Junglas et al., 2020a; Junglas and Schneider, 2018). Thus, rods/tubes and carpets correspond to the predominantly observed Vipp1 structures upon membrane binding. Noteworthy, it has been suggested that Vipp1 carpets are directly involved in membrane protection: upon binding to negatively charged membranes, Vipp1 ring complexes disassemble (Heidrich et al., 2016) and transform into membrane-bound carpets that appear to reduce the proton-flux across damaged membranes (Junglas et al., 2020a). As carpets have only been observed by low-resolution AFM (Junglas et al., 2020a), the detailed molecular structure of the carpets still is elusive.

In this study, we analyzed the structure of Vipp1 in the presence of bacterial membranes using cryo-EM. We observed all previously described species of Vipp1 polymers (ring complexes, stacked-ring assemblies, helical rods, carpets and coats) upon incubating Vipp1 with lipids. The most common species were helical tubes and stacked rings engulfing membranes and membrane-bound carpets. Among these, we solved the cryo-EM structures of three types of helical tubes and four types of stacked ring assemblies at 5 to 7 Å resolution. Apart from helical or ring assemblies, we determined the structures of Vipp1 carpets at three different membrane curvatures to 18 to 20 Å resolution using sub-tomogram averaging. Quantitative analysis of membranes with Vipp1 carpets in electron tomograms suggests that increasing Vipp1 coverage on the membrane correlates with higher local membrane curvature up to a point of complete membrane engulfment in small diameter tubes. By analyzing ten additional unique structures of N-terminally truncated Vipp1 in the presence of lipids, we show that helix α0 is a major determinant for the types of polymeric assemblies that are formed by Vipp1 and that helix α0 serves as a membrane anchor for Vipp1. Moreover, by replacing the Hinge 2 region of Vipp1 with 10 alanines, we generated plasticity-restrained Vipp1 assemblies. The cryo-EM structure of Vipp1 tubes at 3.0 Å resolution revealed the molecular details of the membrane anchoring helix α0 in addition to conserved Vipp1 intersubunit contacts.

## Results

### Vipp1 forms stacked-ring assemblies, helical tubes, carpets, and loose coats upon membrane binding

Recently, the structures of Vipp1 ring complexes were resolved by cryo-EM in the absence of lipid membranes (Gupta et al., 2021; Liu et al., 2021). However, the structure of Vipp1 complexes changes upon membrane interaction, *i.e.,* Vipp1 ring complexes disassemble and transform into carpet structures after/during binding to negatively charged membrane surfaces (Heidrich et al., 2016; Junglas et al., 2020a). To investigate the structure of Vipp1 in the presence of bacterial membranes, we now refolded Vipp1 in the presence of *E. coli* polar lipid extract (EPL). When refolded in the absence of lipids, Vipp1 forms ring complexes and short stacks of rings (**Suppl. Fig. 1A**) in agreement with previous EM observations (Fuhrmann et al., 2009; Gupta et al., 2021; Liu et al., 2021; Saur et al., 2017; Siebenaller et al., 2021). To validate the structural integrity of Vipp1 after refolding, we compared the thermal stability of urea-purified and refolded Vipp1 with natively purified Vipp1, showing that the two samples have essentially identical stabilities (**Suppl. Fig. 1B**). Qualitative assessment of assembly formation as well as thermal unfolding support the notion that refolded protein preparations are structurally similar to the protein purified under native conditions. Upon refolding of Vipp1 in the presence of EPL, we observed the formation of large irregularly shaped vesicles and even maze-like networks of tubulated membranes in contrast to small circularly shaped liposomes in the absence of Vipp1 (**Fig. 1A, Suppl. Fig. 1C**). We scrutinized the Vipp1 cryo-EM micrographs and identified isolated ring complexes as well as elongated rods of stacked rings, tubes, and vesicles covered with carpet structures (**Fig. 1B top**). The observed structures are heterogeneous, *e.g.*, ring complexes have different sizes and the tubular structures are curved, have kinks and variable diameters along the tube axis. When we analyzed the segmented Vipp1 polymer structures by image classification methods, we found 48 % being present as elongated evenly indented rods of stacked rings, 45 % consisting of two types of tubes with distinct helical lattices and 7 % corresponding to regular two-dimensional carpet structures (**Fig. 1B bottom**).

**Figure 1:**
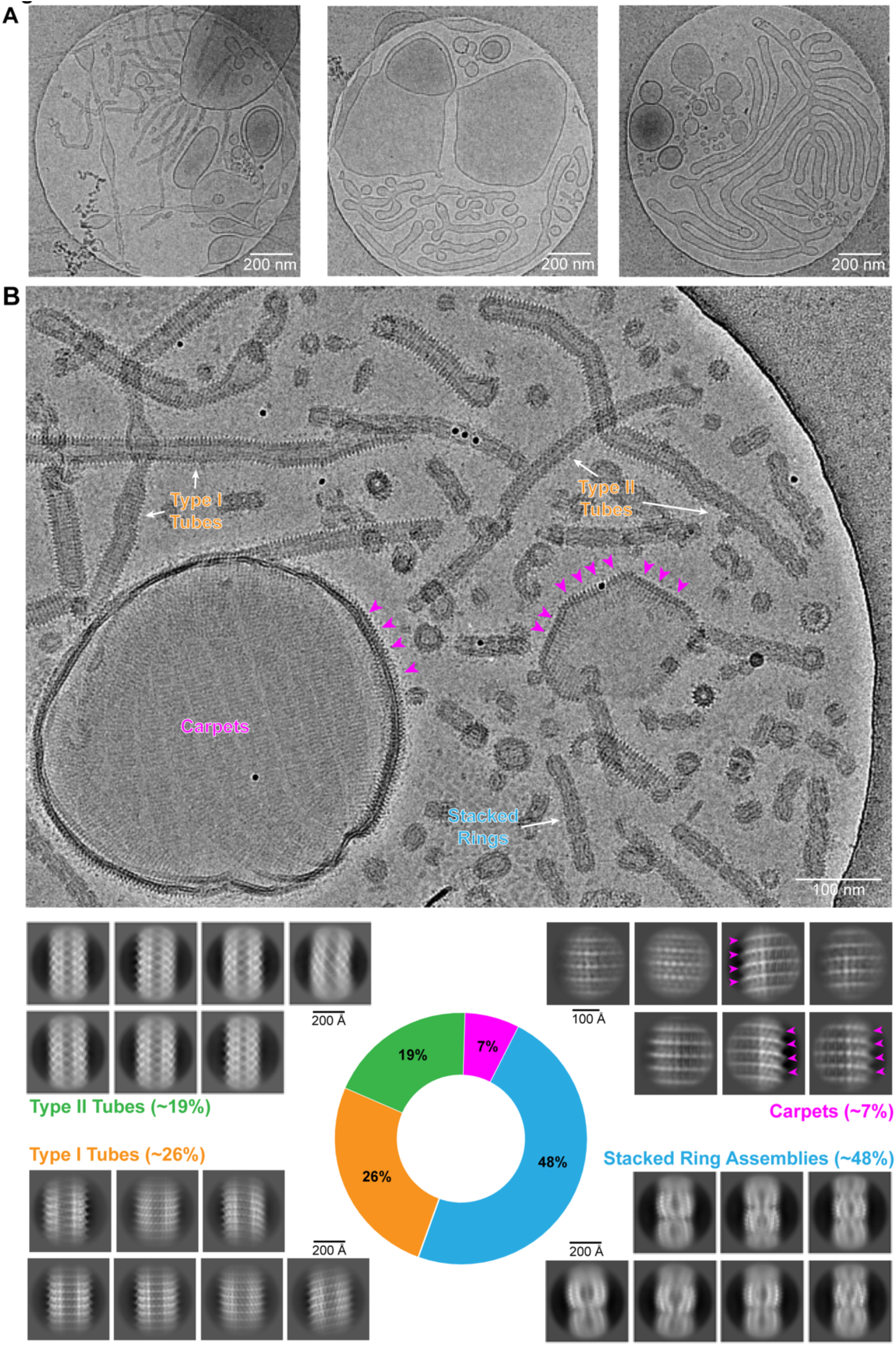
Multitude of different Vipp1 structures after membrane incubation. A: Three overview cryo-micrographs of Vipp1 after reconstitution with EPL lipids. **B:** Cryo-EM micrograph and 2D class averages showing a multitude of different Vipp1 structures after reconstitution with lipids. Apart from ring complexes and stacked-ring assemblies, we identified three additional Vipp1 assemblies: two types of tubular structures with different diameters (Type I and Type II tubes) as well as membrane-bound carpet and loose coat structures. Purple arrowheads indicate pronounced spike-patterns on Vipp1-coated vesicles.

In agreement with previous observations (Gupta et al., 2021), complexes with different diameters and rotational symmetries were identified. The stacked-ring assemblies have a regularly tapered appearance with maximum diameters of individual rings of ∼200 – 280 Å. The stacked-ring assemblies consist of joined end-to-end ring complexes, presumably resulting from stacking of different ring complexes. The Type I tubes displayed a pronounced spike pattern at the outside of the tubes connected by close-to-parallel lattice lines. Most Type I tubes are made of straight and regular stretches of 100 – 200 nm in length with an outermost diameter of 300 Å, interrupted by irregular parts made of bulges, indentations, or kinks. In contrast, Type II tubes were more regular along their length having distinct apparent outer diameters of ∼250 – 260 Å and displayed a crisscross pattern without any pronounced spikes. The carpet assemblies were the most heterogeneous structures. In some cases, they completely covered vesicles and formed ordered almost tape-like tracks on the vesicle surface, while in other cases vesicles were only partially covered by discontinuous patches with pronounced spikes at the edges, closely resembling Type I tubes. Taken together, Vipp1 reconstituted with membranes forms various polymer species, ranging from ring complexes to extended rods, helical rods, carpets, and loose coats.

### Type I and Type II tubes have different monomer orientations with respect to the tube axis

Next, we set out to analyze the structures of the helical Vipp1 tubes. Using single-particle based helical reconstruction, we resolved the structures of Type I tubes at 5.1 Å resolution and two structures of Type II tubes at 7.0 and 7.1 Å resolution, respectively (**Table 1**). Subsequently, we built atomic models of the tubes by flexibly fitting a previously solved monomer structure (PDB:7O3W) into the determined cryo-EM density maps. The Vipp1 secondary structure consists of six α-helices connected by short loops plus a predicted 7^th^ α-helix at the C-terminus (**Fig. 2A** shows the ESCRT-III unified nomenclature for the Vipp1 helices (Schlösser et al., 2023)). In Type I tubes of 305 Å diameter with a helical rise and rotation of 1.79 Å and 124.4°, respectively, the monomers with the characteristic α1-α2 hairpin are arranged almost parallel to the tube axis (**Fig. 2B, Suppl. Fig. 2A and B**). In addition, we found Type II tubes with two different diameters. The smaller tubes had a diameter of 275 Å with a helical rise and rotation of 2.10 Å and 43.5°, respectively. The larger tubes had a diameter of 290 Å with a helical rise and rotation of 1.96 Å and 133.5°, respectively. Apart from the different diameters and helical symmetries, the structures of both Type II tubes were very similar to one another in that the monomers were arranged approximately 45° to the tube axis. Type I and Type II tube cross-sections showed density for an engulfed lipid bilayer in their lumen, with close contact of the Vipp1 monomers to the outer leaflet of the bilayer. The radial density profiles of the helical tubes with enclosed membranes revealed that the phosphate-to-phosphate headgroup distance (from here on bilayer thickness) in all three tubes remained nearly identical at 35 Å (**Fig. 2C**), in agreement with distances previously observed with PspA or CHMP1B (Junglas et al., 2021; Nguyen et al., 2020). The inner leaflet radius changed from 57 Å (Type I tubes) to 41 Å (small Type II tubes).

**Figure 2:**
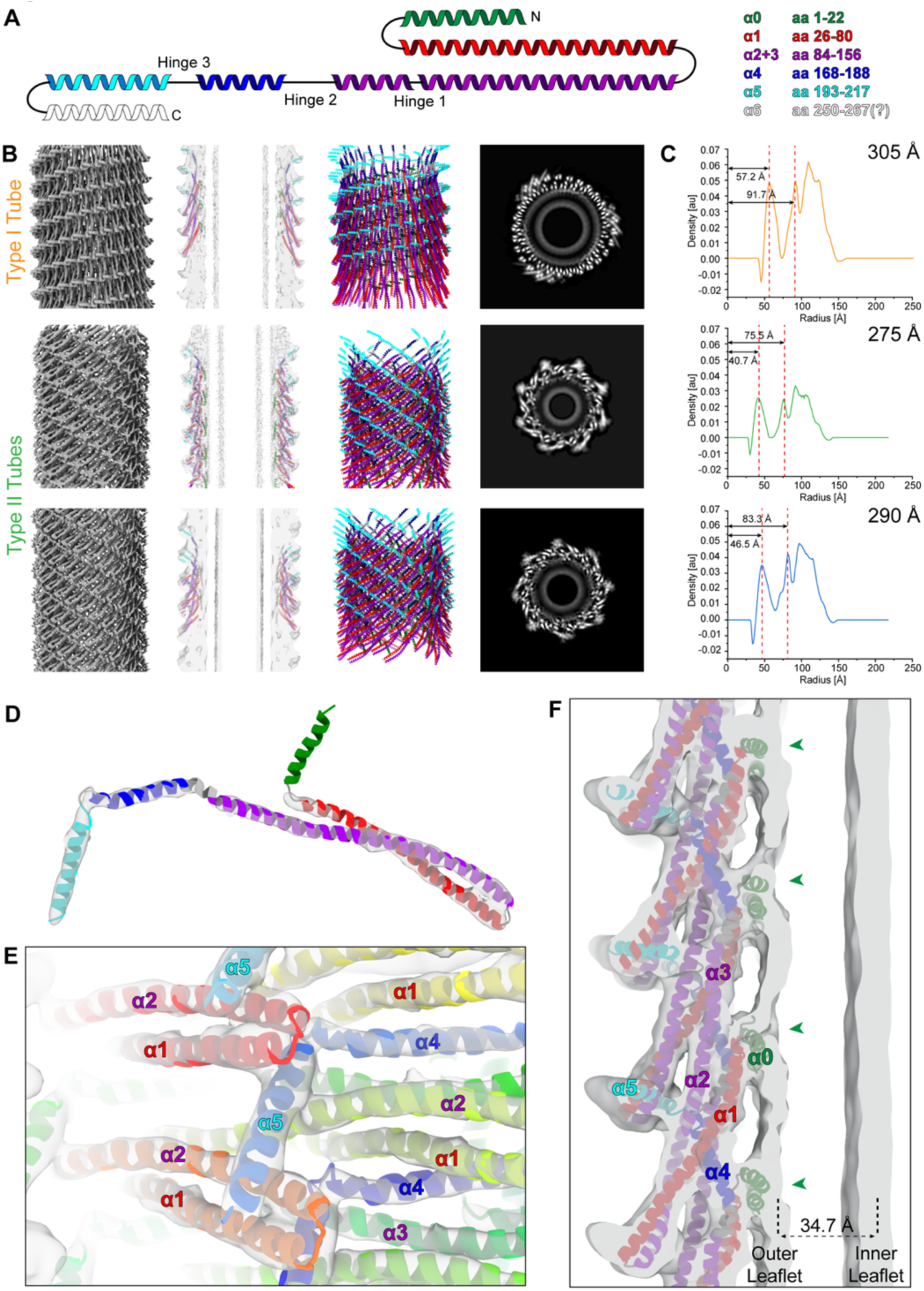
Structural details of Type I and Type II helical Vipp1 assemblies. A: Topology of the Vipp1 structure with color-coded helices: *α*0: green, *α*1 red, *α*2+3 violet, *α*4 blue, *α*5 cyan, *α*6 white. **B:** Cryo-EM structures of Type I (top row) and two Type II assemblies (center and bottom row). Left: cryo-EM maps of Vipp1 tubes. Center left: central xy-slices of cryo-EM maps with the fitted models. Center right: atomic models of Vipp1 tubes in ribbon representation. Right: central z-slices of cryo-EM maps in greyscale. **C:** Radial density profiles of respective Vipp1 tubes (respective outer diameters of the tubes are displayed in the upper right corner of each plot). Dashed lines indicate the peaks of the inner and outer leaflet densities of the tubulated bilayer. **D:** Segmented density and modeled atomic Vipp1 monomer structure found in Type I tubes showing the ESCRT-III fold. **E:** Density of the Vipp1 Type I tubes with the built polymer model showing the polymer core with helices *α*1–4, the tip of the *α*1–3 hairpin, and *α*5 contact sites. **F:** Vipp1 Type I tubes: enlarged view of both leaflets of the tubulated bilayer with the fitted model. Green arrowheads pointing to indentations in the outer leaflet caused by helix *α*0 interacting with the lipids.

**Table 1.**
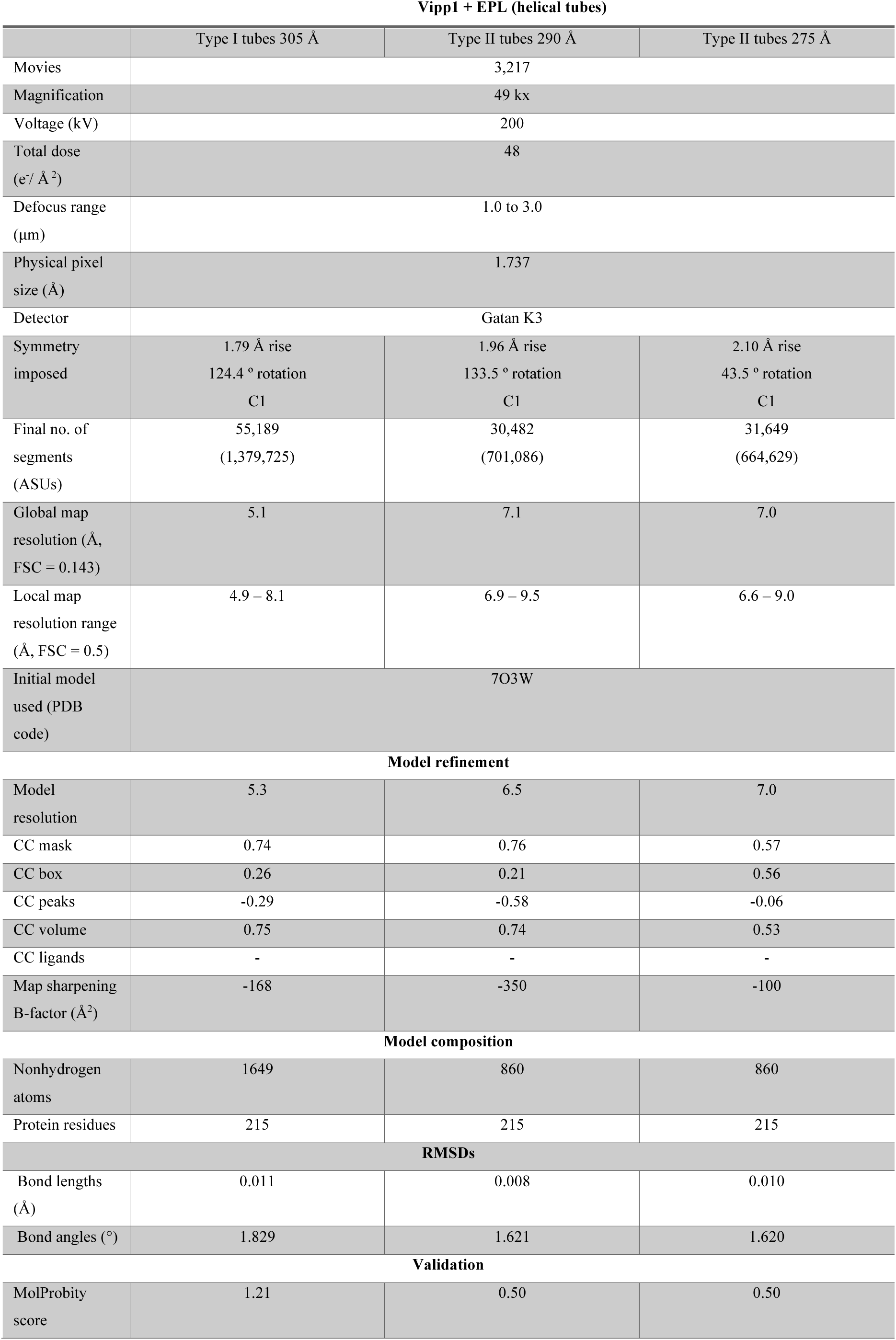
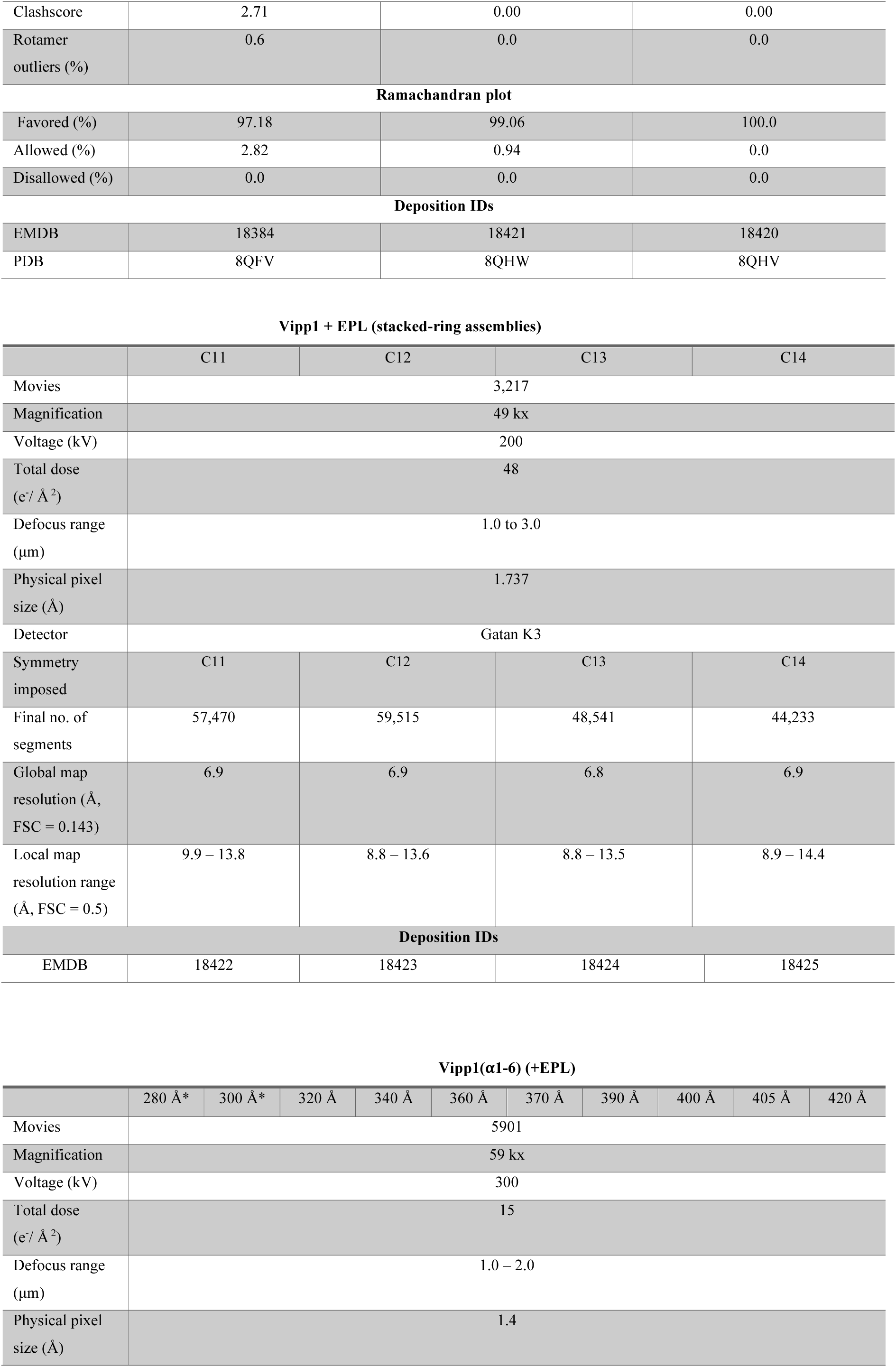
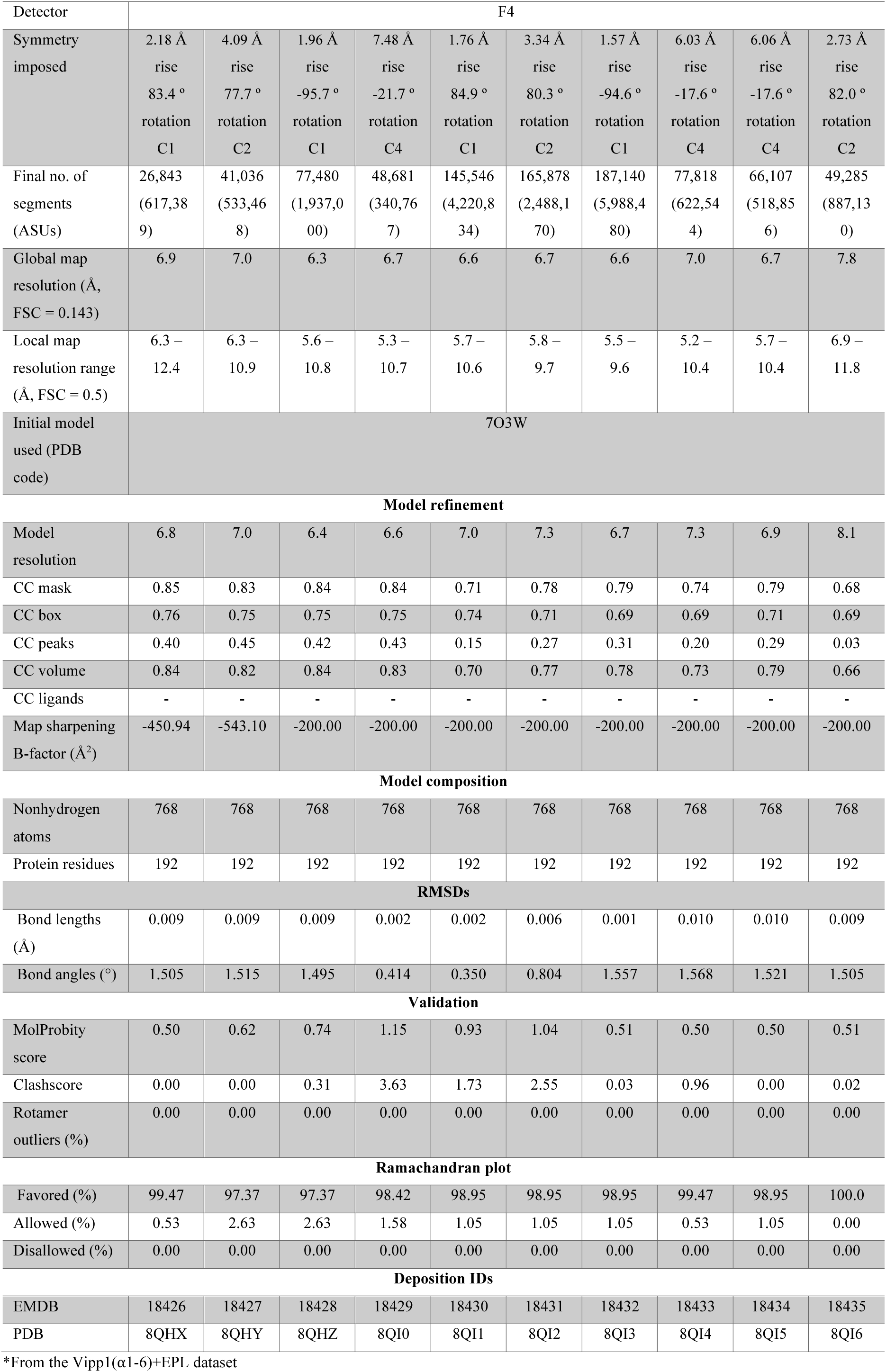
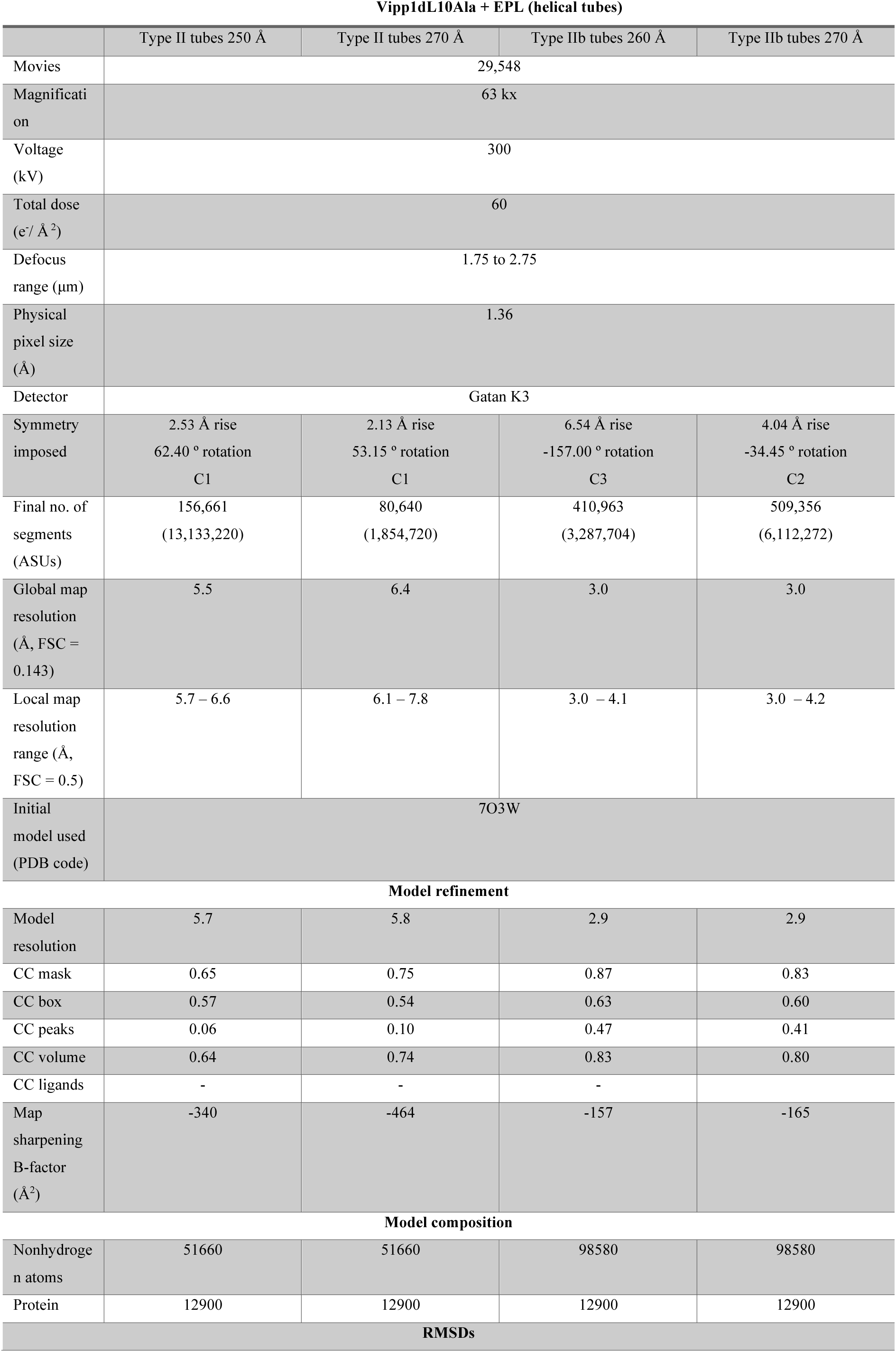
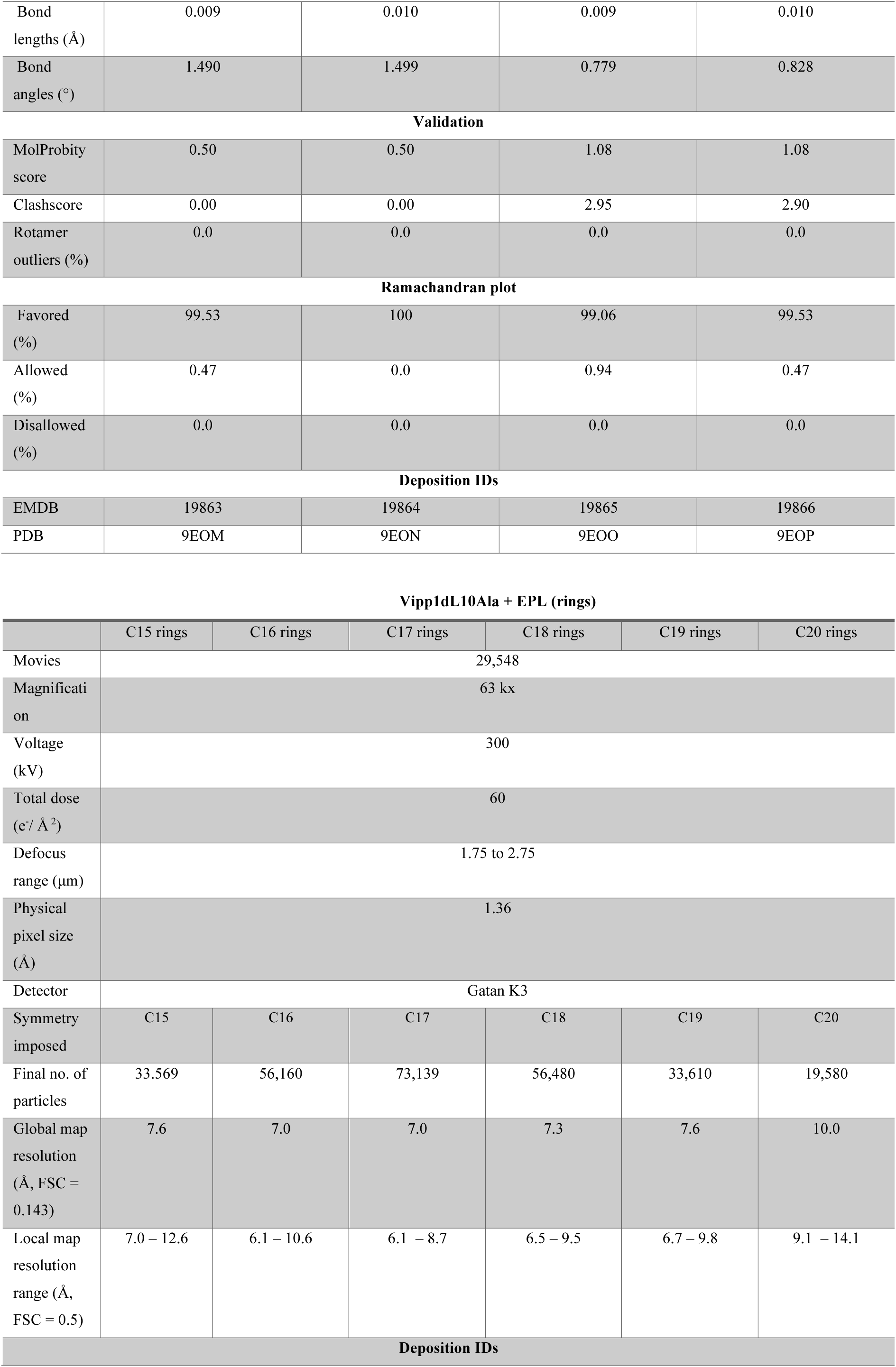
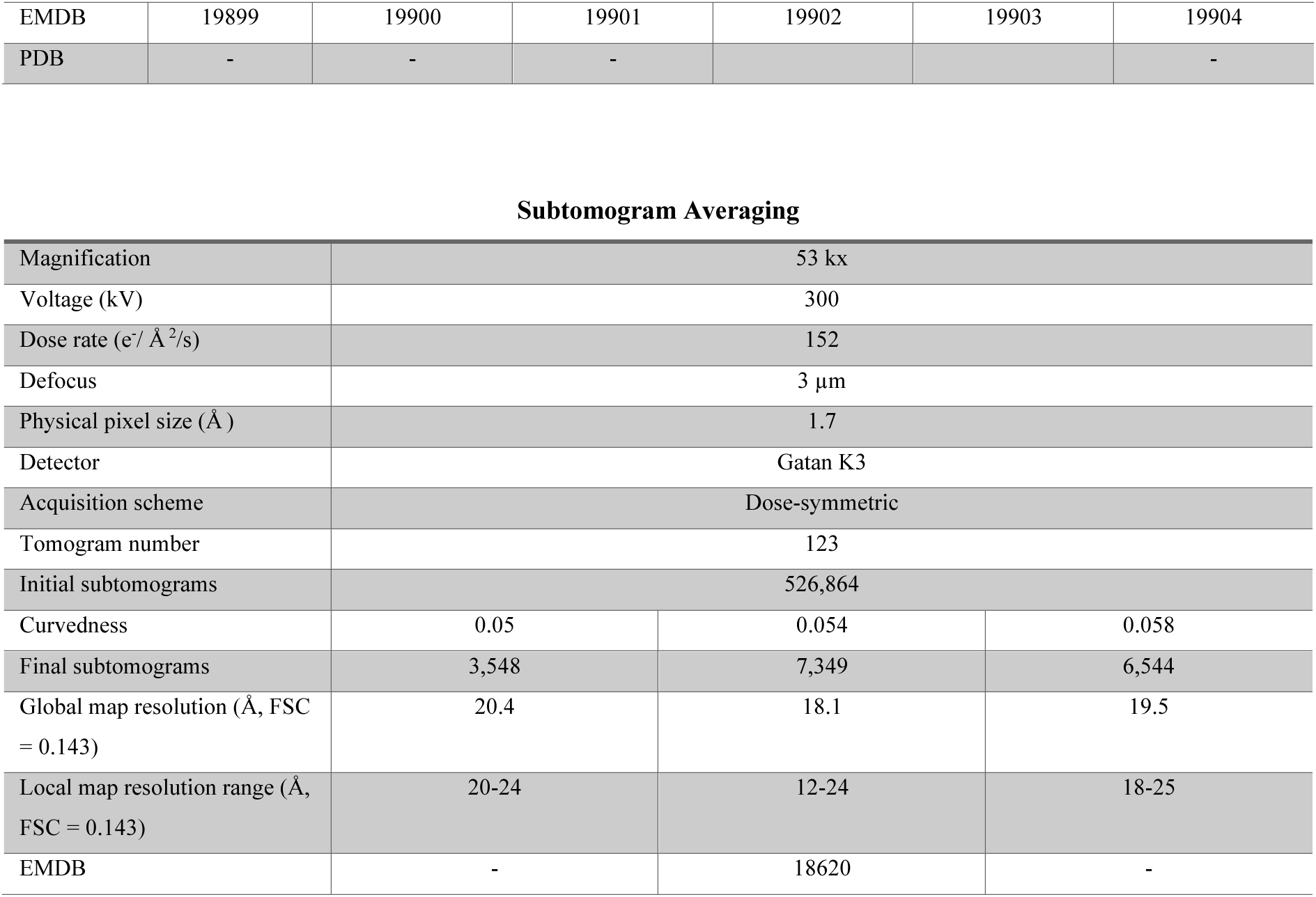
Data collection, image processing, and model refinement Vipp1 assemblies.

In the Type I tube map, we were able to identify α-helical density for α0–5 whereas α6 was not resolved. In the core of the tube wall, the ESCRT-III-typical fold of the helices α1–4 were easily placed in the density (**Fig. 2D**). In the periphery, the tip of the α1–3 hairpin and α5 contact sites create the distinctive structural motif when seen in the side view (**Fig. 2E**). Although the polymeric assemblies differ in their architecture, the intermolecular contacts that stabilize the assemblies are nearly identical in Type I and Type II tubes (α1–4 contacts in the core, α1–3 hairpin-to-α5 contacts in the periphery). These or similar intermolecular contacts are conserved in Vipp1 rings, PspA rods, and eukaryotic ESCRT-III proteins (Junglas et al., 2021; Liu et al., 2021; Schlösser et al., 2023). To accommodate similar contacts in different topologies, the monomers adapt with changes at the flexible hinge regions 1, 2, and 3 (**Suppl. Fig. 2C**). Within the Vipp1 Type I and Type II tubes, membrane contact is mediated by helix α0 that is lying flat in the membrane plane causing a local disturbance in the membrane, presumably by pulling the lipid headgroups towards the inner tube wall (**Fig. 2F, Suppl. Fig. 2D**). Helix α0 appears to partially submerge into the outer leaflet of the bilayer. In summary, Vipp1 helical tubes display two distinct monomer topologies with respect to the tube/membrane axis while the molecular contacts between the monomers or of the monomers with the membrane remain essentially identical between the different assemblies: helix α1–4 contacts stabilize the core, α1–3 hairpin-to-α5 contacts stabilize the periphery, and helix α0 mediates interaction with the outer leaflet of the membrane.

### Joined Vipp1 ring assemblies induce high local membrane curvature

In the above-presented micrograph, we identified end-to-end joined ring complexes, *i.e.*, stacked-ring assemblies, with different rotational symmetries (see **Fig. 1B**). Based on the cryo-EM images, we determined the 3D structures of C11 to C14 stacked ring assemblies at resolutions of 6.8 to 6.9 Å (**Fig. 3A**, **Table 1**). The individual ring complexes in these assemblies had C11 – C14 rotational symmetry. Although we also found ring complexes with higher/lower rotational symmetries, we were not able to generate reliable 3D reconstructions in those cases. Subsequently, we flexibly fitted models of the available Vipp1 ring structures (Gupta et al., 2021; Liu et al., 2021) into our density maps, and found that the available models had good overlap with our reconstructions. Only minor adjustments at hinge 3 and helix α5 were necessary, presumably reflecting the poorer resolution and map quality of the stacked ring assemblies compared with the single ring complex structures (EMD 11468 (C11), 11469 (C12), 11470 (C13), 12710 (C14), PDB 6ZVR (C11), 6ZVS (C12), 6ZVT (C13), 7O3W (C14)) (**Fig. 3A, Suppl. Fig. 3A**). In those elongated structures, the rings are joined end to end resulting in a polar head-to-tail stacking: The more tapered side of one ring assembly (*i.e.,* the top side) is connected to the less tapered side (*i.e.,* the bottom side) of the next ring assembly. Unlike the individual ring complexes, our stacked-ring assembly structures show bilayer density in the lumen with the inner membrane leaflet clearly discernable. As the outer leaflet is located close to the denser ring wall, it is not easily distinguished from the ring density. In analogy to the Type I and Type II helical tubes, helix α0 is lying flat on the membrane and appears to partially submerge into the outer leaflet (**Fig. 3B**). Due to the tapered nature of the ring complexes, the membrane diameter changes over the z-axis of a ring. In the C12 rings, the inner leaflet diameter changes in accordance with the Vipp1 structures from 42 to 86 Å from the narrowest to the widest part of the ring (**Fig. 3C**). We observed similar effects for the other ring assemblies, and, as expected, the ring complexes with the smallest diameters engulf the membranes with the smallest diameters (**Suppl. Fig. 3B**). Interestingly, our maps include density of the adjacent upper and lower neighboring ring stacks. The contact between the ring complexes is mediated by the interaction of the α1–3 hairpin of the lower ring to α5 of the upper ring complex. Additionally, the stacked ring assemblies include a continuous membrane tube engulfed by the stacked ring assembly. The engulfed membrane is most constricted at the position where two rings touch, further increasing the local curvature (**Fig. 3D, Suppl. Fig. 3C**). In contrast to the thickness of the lipid bilayer engulfed by helical Vipp1 tubes, the bilayer thickness in stacked Vipp1 rings was generally smaller and more variable (**Suppl. Fig. 3D**). First, the bilayer thickness within one ring varies from the narrowest to the widest part (*i.e.,* 32 to 34 Å in the C12 rings). Second, the bilayer thickness varied from the smaller to the larger rings (*i.e,.* 32 to 30 Å at the narrowest part of C11 to C14 rings). In summary, stacked Vipp1 ring assemblies appear to have the same structure as individual ring complexes, with the monomer structures and hairpin angles relative to the membrane axis changing in each layer from the bottom to the top layer of each ring. Additionally, stacked ring assemblies engulf a continuous lipid bilayer with high local curvature at the contact site of two rings.

**Figure 3:**
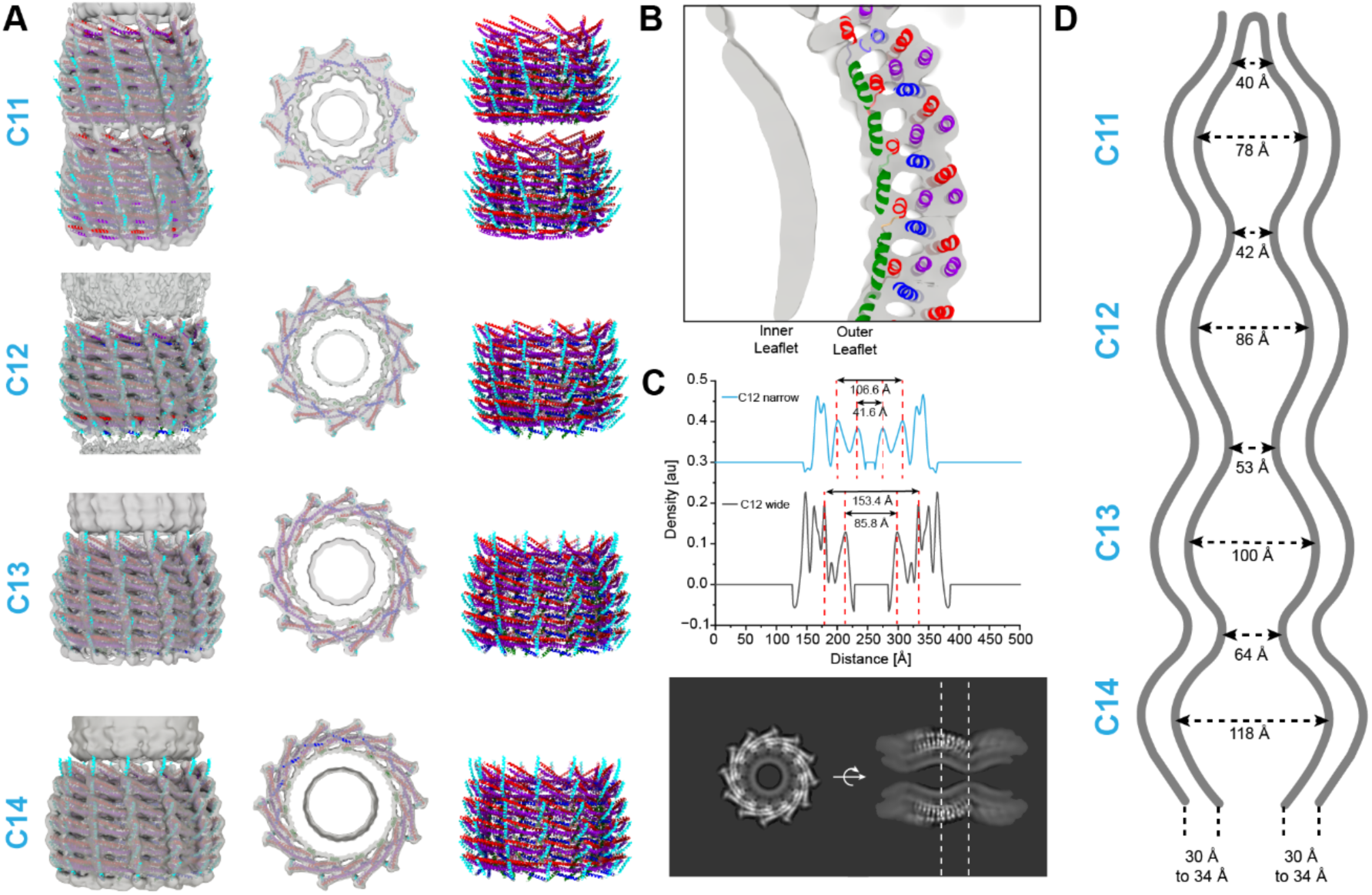
Cryo-EM structures of stacked Vipp1 ring assemblies with engulfed membranes. A: Structures of stacked Vipp1 ring assembly with C11, C12, C13, and C14 rotational symmetry (top to bottom rows, respectively). Left: transparent cryo-EM maps with the fitted models. Center: central z-slices of cryo-EM maps with fitted models. Right: atomic models of Vipp1 rings in ribbon representation (*α*0: green, *α*1 red, *α*2+3 violet, *α*4 blue, *α*5 cyan). **B:** Enlarged view of a Vipp1 ring in a stacked-ring assembly showing both leaflets of the tubulated bilayer with the fitted model. **C:** Top: Density profiles across the stacked C12 ring complex. Bottom: z (left) and xy-slice (right) of the map in greyscale including the locations of the density sections above (dashed lines). **D:** Schematic sketch of the measured membrane distances in stacked Vipp1 ring assemblies for the narrowest and widest part of C11 – C14 rings and the observed bilayer thicknesses (in detail shown in **Suppl.** Fig. 3B**-D**).

### Helix α0 is critical for membrane tubulation and affects the Vipp1 polymer structure

The here-determined structures of Vipp1 polymers in the presence of membranes indicated that helix α0 is immersed in the outer lipid leaflet of the bilayer. To obtain more insights into its function, we analyzed the interaction of a truncated Vipp1 lacking helix α0 (Vipp1(α1-6)) with membranes using cryo-EM. In the absence of membranes, Vipp1(α1-6) formed large and straight rods of different diameters (**Suppl. Fig. 4A**). In contrast to the full-length Vipp1, we did not observe the formation of any ring complexes nor any other polymers than Vipp1 rods, in agreement with previous observations (Thurotte and Schneider, 2019). After reconstitution with lipids, Vipp1(α1-6) formed again solely rods, similar to the rods observed in the absence of membranes. Additionally, in the micrographs we did not find any indications of Vipp1(α1-6) binding to membrane surfaces, neither protein carpets, nor loose coats on the membranes, nor deformed/irregularly shaped vesicles (**Suppl. Fig. 4B**). The Vipp1(α1-6) rods had a large range of diameters from 280 to 420 Å according to the class averages and the radial density profiles of the helical reconstructions (**Suppl. Fig. 4A-C**). We also compared the radial density profiles of Vipp1(α1-6) rods with full-length Vipp1 tubes in the presence of membranes (**Suppl. Fig. 4D**). Strikingly, we could not find density for a tubulated membrane in the Vipp1(α1-6) rods. Therefore, we conclude that helix α0 is critical for membrane tubulation and engulfment by Vipp1 polymers. In total, we determined a series of ten unique rod structures with global resolutions of 6.3 to 7.8 Å, respectively, with the best-resolved parts in the center of the rod walls and poorer resolutions in the periphery (**Suppl. Fig. 4E and F**). The Vipp1(α1-6) structures obtained in the absence and presence of lipids were mostly identical in the respective rods including the same helical symmetries. Again, we used PDB: 7O3W as the reference monomer structure and flexibly fitted it to the rod reconstructions. We noticed that the monomer arrangement was different from our full-length Vipp1 helical tubes and was more similar to *Syn*PspA rods (*i.e.,* the angle of the monomers was ∼90 ° relative to the rod axis) (Junglas et al., 2021). However, the before-mentioned features of Vipp1 polymers remained conserved (α1–4 contacts in the core, α1–3 hairpin-to-α5 contacts in the periphery), and, as for the other Vipp1 assemblies, the monomers displayed a structural plasticity due to the conformational flexibility in the hinge regions 1-3 to adapt to the different rod diameters (**Suppl. Fig. 5A and B**). In conclusion, helix α0 is critical for membrane interaction and tubulation of Vipp1 as well as in controlling the polymer architecture of Vipp1.

### Hinge 2 is critical for the structural plasticity of Vipp1 assemblies

The here determined Vipp1 structures in the presence of lipids reveal a remarkable degree of structural plasticity (see **Fig. 1B**). Mainly responsible for the associated conformational flexibility appears to be the loop region between helix α3 and α4 (Hinge 2) that enables different shapes and extensions of the Vipp1 monomers to adapt to different ring and rod diameters, *e.g.*, as demonstrated for the Vipp1(α1-6) rods (see **Suppl. Fig. 5B**) and Vipp1 rings (Gupta et al., 2021; Liu et al., 2021). Thus, we hypothesized that limiting the conformational flexibility of this region in the Vipp1 monomer may reduce the plasticity of Vipp1 assemblies observed during membrane interaction. For this purpose, we designed two Hinge 2 mutants (**Fig. 4A**). First, we created a mutant where the whole loop connecting α3 and α4 was removed (aa 157 – 166, Vipp1 dL10). Unfortunately, this mutant did not form any well-ordered assemblies suitable for more detailed structural characterization, suggesting that Hinge 2 is indeed critical for Vipp1 assembly formation (**Suppl. Fig. 6A left**). Second, we replaced the loop region with a deca-alanine stretch (Vipp1 dL10Ala) to convert Hinge 2 to an α-helix, thus potentially stabilizing the conformation of large diameters while also limiting the structural flexibility of the loop. Interestingly, this mutant was capable of forming single-ring complexes in the absence of membranes, although the ring complexes had larger diameters and were not as uniformly shaped as the wildtype (WT) ring complexes (**Suppl. Fig. 6A right** and compare **Suppl. Fig. 1A)**. Next, we analyzed the structure of Vipp1 dL10Ala in the presence of membranes using cryo-EM, as we did for the WT. We found that Vipp1 dL10Ala formed mostly formed Type II tubes (69%) including single-ring complexes (27%), stacked-ring assemblies (3%) as well as Type I tubes (2%) (**Fig. 4B, Suppl. Fig. 6B**). In contrast to WT, we found only minor shares of two-dimensional carpet or stacked-ring structures and the observed tubular structures were straight in appearance without any kinks and bends but with a constant apparent diameter of 220 Å along the tube axis. Thus, the spectrum of assemblies found in the Vipp1dL10Ala EPL sample was practically reduced to two types of assemblies (Type II tubes and single ring complexes) making up approx. 95% of all assembly types. In contrast, in the WT EPL sample, all five assembly types were more or less evenly distributed. These data indicate that the replacement of Hinge 2 with 10 alanines indeed constrained the observed plasticity of Vipp1 assemblies.

**Figure 4:**
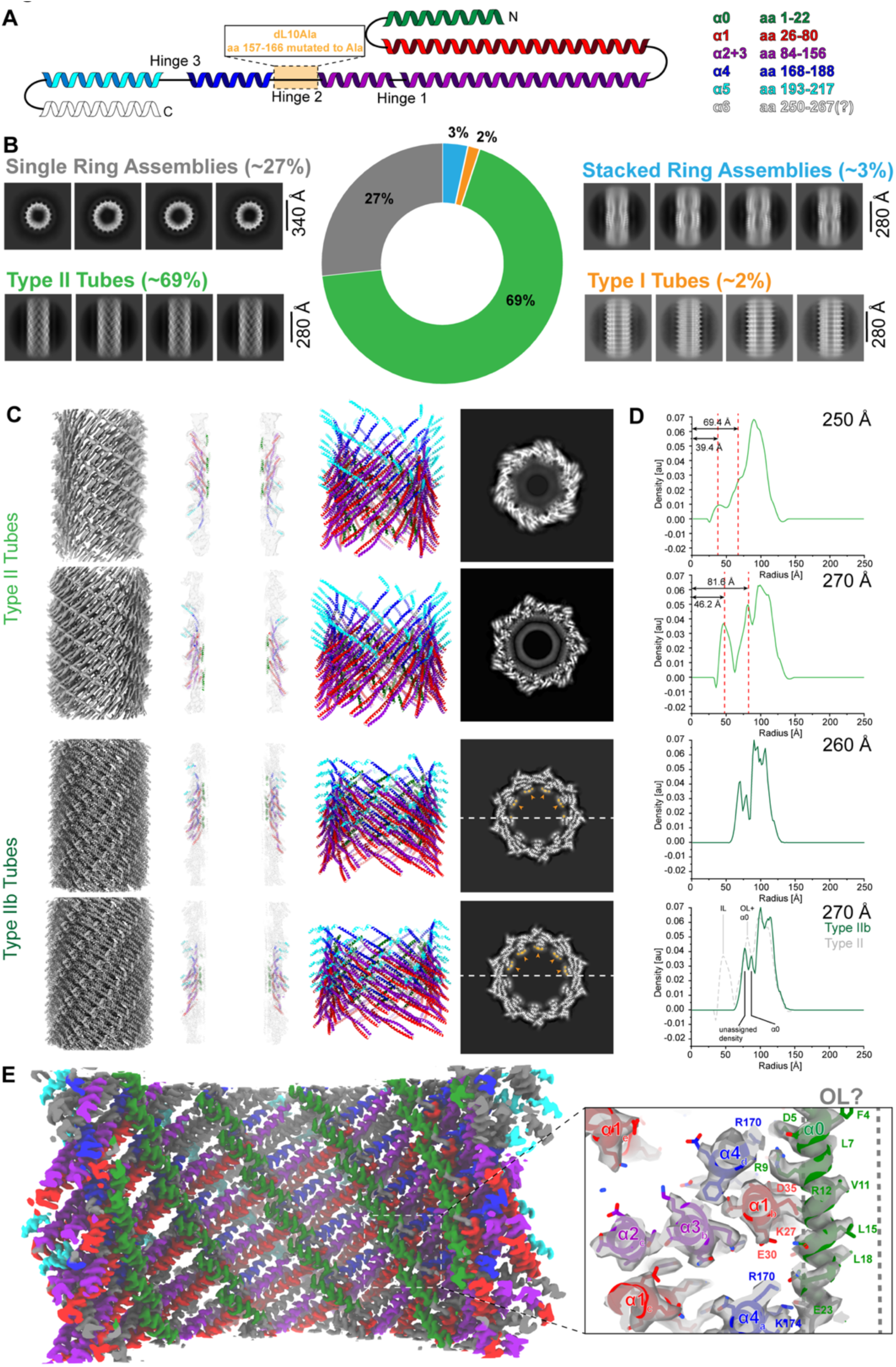
Cryo-EM structures of the plasticity-reduced Vipp1 dL10Ala. A: Topology of the Vipp1 structure with indicated mutation site for Vipp1 dL10Ala and color-coded helices: *α*0 green, *α*1 red, *α*2+3 violet, *α*4 blue, *α*5 cyan, *α*6 white. **B:** 2D class averages showing different Vipp1 dL10Ala structures after reconstitution with lipids. Apart from ring complexes and stacked-ring assemblies, we identified Type I and Type II tubes. **C:** Cryo-EM structures of two Type II (top rows) and two Type IIb assemblies (bottom rows). Left: cryo-EM maps of Vipp1 dL10Ala tubes. Center left: central xy-slices of cryo-EM maps with the fitted models. Center right: atomic models of Vipp1 tubes in ribbon representation. Right: central z-slices of cryo-EM maps in greyscale with highlighted unassigned density in orange. **D:** Radial density profiles of respective Vipp1 dL10Ala tubes (respective outer diameters of the tubes are displayed in the upper right corner of each plot). Dashed lines indicate the peaks of the inner and outer leaflet densities of the tubulated bilayer. Bottom profile shows the superposition of Type IIb (solid green) and Type II (grey dashed) tubes of 270 Å diameter assigned with putative structural elements. **E:** Cryo-EM map of Vipp1 dL10Ala Type IIb tubes with surface colored Vipp1 *α*-helices and enlarged view of the cryo-EM map with the fitted model showing the helix *α*0 interactions and putative position of a membrane outer leaflet (OL).

Given the reduced plasticity of the Vipp1dL10Ala assemblies, we set out to determine the cryo-EM structures from this sample. First, we focused on single-ring complexes and were able to determine six structures ranging from C15 to C20 symmetry (**Suppl. Fig. 6C)**. As observed in negative stain, the single-ring complexes were larger than the WT ring complexes and showed C15 to C20 rotational symmetry with diameters from 330 to 435 Å, while the so far solved WT ring complex structures have C11 to C18 symmetry with approx. 240 to 350 Å diameters (Gupta et al., 2021; Liu et al., 2021). The resolution of our single-ring complex reconstructions was limited to 7 to 10 Å resolution, presumably due to a limited homogeneity (**Suppl. Fig. 6D**). Due to the small fraction of stacked ring assemblies and Type I tubes present in the Vipp1 dL10Ala sample, we were not able to generate reliable reconstructions for these assemblies. As for the WT, we identified two sizes of regular Type II tubes with an engulfed lipid bilayer. The smaller tubes had a diameter of 250 Å with a helical rise and rotation of 2.53 Å and 62.4°, respectively. The larger tubes had a diameter of 270 Å with a helical rise and rotation of 2.13 Å and 53.15°, respectively. Apart from the different diameters and helical symmetries, the structures of both Type II tubes were similar to the WT Type II tubes (the monomers were arranged approximately 45° to the tube axis, and the engulfed membrane tubes had inner leaflet radii of 39 and 47 Å, respectively) (**Table 1**, **Fig. 4C and D**). The regular Type II tubes were resolved at 5.5 and 6.4 Å resolution, respectively (**Suppl. Fig. 7A**). In addition to the regular Type II tubes, we found a subclass of Type II tubes making up approx. 46% of the overall sample, apart from 23 % regular Type II tubes. These Type IIb tubes formed left-handed instead of right-handed helices and also came in two sizes. The smaller tubes had a diameter of 260 Å with a helical rise and rotation of 6.54 Å and -157.0° in addition to C3 symmetry, respectively. The larger tubes had a diameter of 270 Å with a helical rise and rotation of 4.04 Å and -34.45° in addition to C2 symmetry, respectively. Apart from the different diameters and helical symmetries, the structures of both Type II tubes were similar to the WT Type II tubes with the monomers arranged approximately 45° to the tube axis. Both Type IIb tubes were resolved at 3.0 Å resolution, presumably due to the reduced flexibility of the Hinge 2 region (**Suppl. Fig. 7A**), allowing model building at near-atomic resolution.

Comparing the monomer structures of the mutant with the WT reveals high similarity in the overall monomer architecture. However, the dL10Ala monomer models had an overall extended length and showed less flexibility in the Hinge 2 region, especially in single-ring complex monomers (**Suppl. Fig. 7B**), which also explains the observed larger diameters of dL10Ala rings, as smaller rings require stronger bended and less extended monomers. The Type II tubes have one continuous approx. 170 Å long α-helix formed by helix α2, α3, and α4 presumably stabilized by the introduced deca-alanine stretch (**Suppl. Fig. 7C**). The monomer structure of the Type IIb tubes revealed another unique feature of this assembly, namely the fractured structure helix α5 that is split into two separate helices while still maintaining the canoncical ESCRT-III α1–3 hairpin-to-α5 contacts (**Suppl. Fig. 7D**). The built Type IIb model also revealed the detailed structure of Hinge 2 in Vipp1 dL10Ala (**Suppl. Fig. 7E**). As predicted, the flexible loop was replaced by a small kink at A163 and accordingly helix α3 as well as α4 were elongated, making Hinge 2 less flexible. Similarly, the inter and intramolecular interactions within the tube were resolved at near-atomic detail. The intramolecular interaction between helix α1 and α2 is primarily based on hydrophobic interactions, *e.g.,* L108, Y104, L97, T65, L61, T57, and I51 (**Suppl. Fig. 7F**). As shown previously, a critical intermolecular interaction to stabilize higher-order Vipp1 assemblies involves the conserved _168_FERM_171_ motif (Heidrich et al., 2016; Junglas et al., 2020a; Saur et al., 2017). F168 interacts with a hydrophobic groove (L29, V33, A132, K135, and L139) between helix α1 and α3 of an adjacent monomer (**Suppl. Fig. 7G**). E169 is directed into a gap between four neighboring monomers and has no direct interaction partners. R170 and M171 point to the lumen of the tubes and interact with helix α0 of two other subunits (R6 (subunit a), and E23, P25 (subunit b)). Another conserved residue close to the _168_FERM_171_ motif is E179, interacting with R142 in α3 of an adjacent subunit.

As shown in this study (see **Suppl. Fig. 4 and 5**), helix α0 was found to be critical for membrane tubulation while it also affects the Vipp1 polymer structure. Given its key position at the inner wall of the Vipp1 assembly, it contacts a total of four different subunits in Vipp1 assemblies by forming polar interactions via its own residues D5, R6, R9, R12, and E23 with subunit (a) α4 R174, subunit (b) helix α0 N16, E23 and α1 D35, subunit (d) helix α4 R170, and subunit (f) helix α0 R6 (**Fig. 4E**), which may explain the central importance of helix α0 for enabling different assembly states of Vipp1, *i.e.*, rings, Type I/II Tubes *vs*. solely rods in the Vipp1 (α1-6) sample. In the Type IIb tubes, helix α0 is not embedded in a continuous bilayer as in the regular Type II tubes. Instead, we found unassigned discontinuous density in the tube lumen, at a radius of 73 or 81 Å, respectively, at the similar radii where the head group peak of the bilayer of regular Type II tubes was found (see **Fig. 4C** highlighted in orange, **Fig. 4D** bottom overlay Type II and Type IIb). In line with previous observations (Gupta et al., 2021), we propose that this density is caused by individual lipid molecules interacting with helix α0. The orientation of helix α0 with respect to the α1/α2 hairpin is identical in the Type IIb tubes in comparison with Type II and all previously described Vipp1 assemblies (single/stacked ring assemblies, Type I tubes (**Fig. 4E left** compare with **Suppl. Fig. 2C and Suppl. Fig. 7B-D**)). Moreover, as the radial position of helix α0 is identical at approx. 84 Å as well as helices α1/2 for bilayer-internalized Type II and membrane-free Type IIb assemblies, we will use the better resolved 3.0 Å structure of Type IIb assemblies for further side-chain based interpretations.

**Figure 5:**
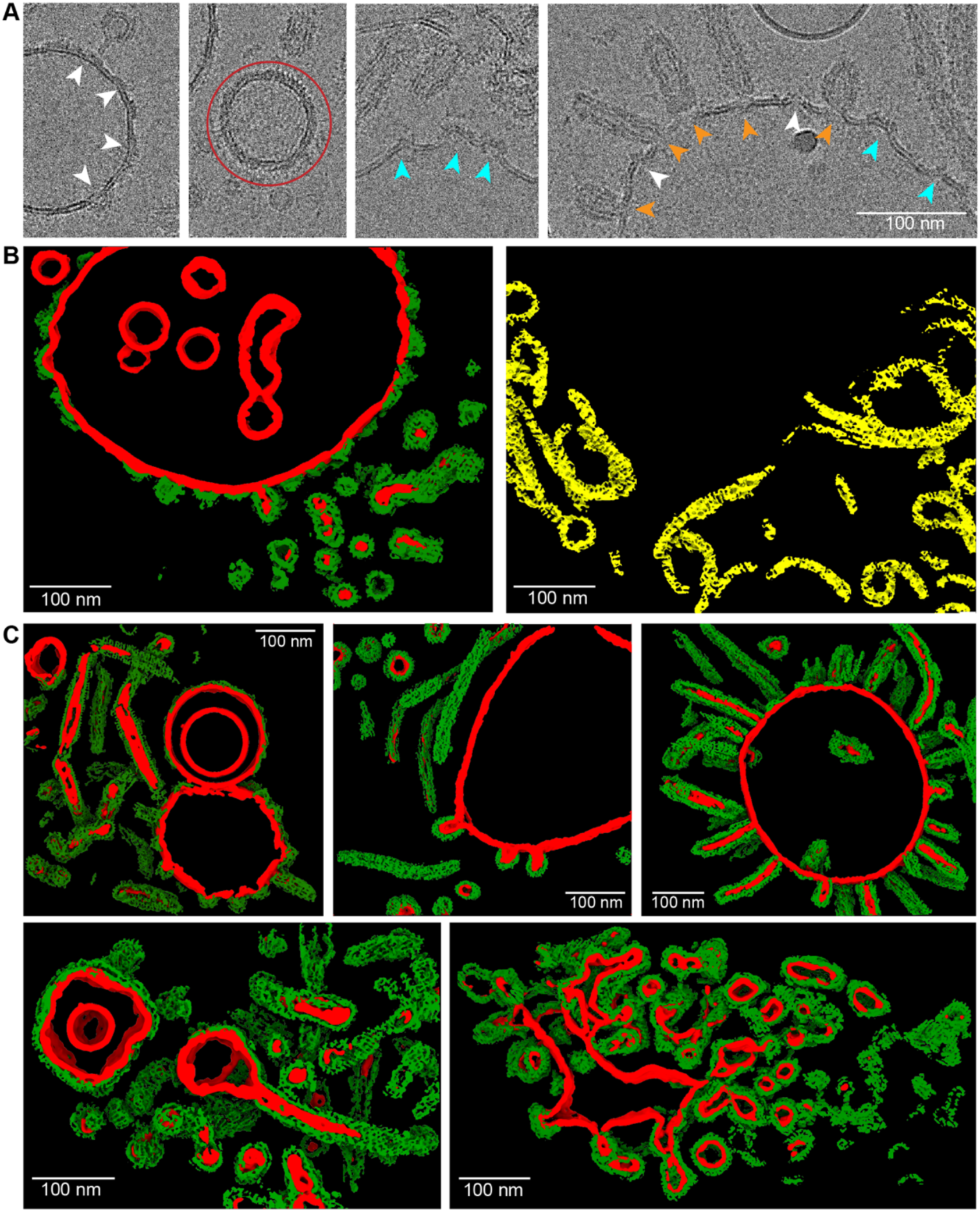
Assemblies of Vipp1 loose coats and carpets on membranes. A: Cryo-EM micrographs of Vipp1 after lipid reconstitution with EPL. White arrowheads indicate loose coats of Vipp1 on flat membranes. Cyan arrowheads indicate sites of small Vipp1 patches with induced bulges at the membranes. Orange arrowheads indicate sites where Vipp1 ring or rod assemblies bind to the membrane or bud off the membrane. The red circle indicates a vesicle fully covered with a Vipp1 carpet. **B:** 3D renderings of the same tomogram after segmentation using a progressively trained U-Net. The U-Net was trained to segment membranes, membrane coverings, and spirals into separate classes. For better visibility, membranes, Vipp1, and Vipp1 spiraling ribbons were rendered in red, green (left), and yellow (right), respectively. The source raw tomogram is shown in **Suppl. Fig. 8C**. Red: membrane, green: Vipp1 rings, rods, and carpets, yellow: Vipp1 spirals. **C:** Gallery of segmented Vipp1 lipid tomograms.

In the Type IIb structure, we found that the side chains L3, F4, L7, V10, V11, L15, L18, and V19 making up a hydrophobic face of helix α0 are directed into the inner tube lumen. They interface with additional density supporting the notion of interacting with hydrophobic acyl chains of lipids, respectively. The opposite face of helix α0, directed to the Vipp1 wall pointing outwards of the assembly, is lined by positively charged residues R6, R9, and R12 interfacing with negatively charged residues, *e.g.*, D35 (helix α1) while they could possibly also interact with negatively charged lipid-head groups (**Fig. 4E right, Suppl. Fig. 7H**). To experimentally test which helix α0 residues are involved in membrane interaction, we made use of the observation that the isolated Vipp1 helix α0 forms an α-helix only when binding to negatively charged membrane surfaces (McDonald et al., 2017). In this way, we measured the membrane binding capacity of different variants of the helix α0 peptide using CD spectroscopy. Compared with the WT peptide, peptides that had R6 and R9 or F4 and V11 replaced by two alanines reduced the membrane affinity to a small extent as they reached similar ellipticity plateaus, while the WT peptide binding curve decayed faster than the R6AR9A and F4AV11A binding curves (**Suppl. Fig. 7I**). By replacing the eight hydrophobic residues with alanines, thus destroying the amphipathic character of the peptide, we abolished membrane binding completely. This observation indicates that the amphipathic character is essential for membrane interaction while the hydrophobic face appears to be more critical than the positively charged residues. In conclusion, we could show that Hinge 2 is a critical element for the structural plasticity of Vipp1 assemblies. Using the plasticity-reduced Vipp1 dL10Ala variant, we successfully determined the structure of Vipp1 tubes at near-atomic resolution and revealed critical interactions of helix α0 important for the membrane/lipid interaction as well as for the intermolecular assembly contacts.

### Vipp1 membrane coverage determines membrane curvature

In addition to helical tubes, stacked-ring assemblies, and ring complexes, we also found carpets and loose coats of Vipp1 on membranes in the cryo-EM micrographs of the Vipp1 lipid sample (see **Fig. 1B**). When taking a closer look at the small Vipp1 coats, we found discontinuous densities on the membrane of vesicles without changing the local curvature (**Fig. 5A** white arrowheads). However, when patches exceeded a critical size, they were found on positive local curvature on the vesicular surface introducing noticeable bulges emanating from the membrane plane (**Fig. 5A** cyan arrowheads). When membrane-attached ring or stacked-ring assemblies were observed, they also showed local curvature formation at the host membrane (**Fig. 5A** orange arrowheads). In many cases, the curved membrane extended to a small neck connected to the ring complexes or longer stacked-ring assemblies. Other discernible species of membrane-bound Vipp1 assemblies were carpets that fully cover vesicles (**Fig. 5A** red circle). These vesicles typically had a smooth spherical shape without bumps or bulges while they exhibited a pronounced spike pattern at the edges and a regular stripe pattern in the center. A comparison of the cylindrical projection of the Vipp1 Type I tubes and 2D class averages of the carpet and their corresponding Fourier transforms revealed that the distances occurring in this regular stripe and spike pattern were very similar, indicating common underlying structures (**Suppl. Fig. 8A and B**). These distances of the cylindrical projection in Type I tubes correspond to the stacking distance of the α1/α2 helical hairpin array suggesting that a similar hairpin array is also present in Vipp1 carpets.

As the membrane-attached Vipp1 assemblies were not suitable for 3D reconstruction from 2D images, due to limited views of preferred orientation, we elucidated the 3D structure of these assemblies by cryo-electron tomography (cryo-ET). Therefore, we collected a total of 123 tomograms from a Vipp1 lipid sample suitable for further analysis. Subsequent membrane segmentation using a progressively trained U-Net (Heebner et al., 2022) revealed a large vesicle with several smaller encapsulated vesicles (**Fig. 5B** left, raw tomogram in **Suppl. Fig 8C**) while the large vesicle is covered by Vipp1 assemblies, such as loose coats and larger carpets. Noteworthy, we also discovered tape-like spiraling ribbons of Vipp1 in our tomograms (**Fig. 5B** right, **Suppl. Fig. 8D**). These highly curved 2D assemblies were only 1-2 nm thick and had lengths of several hundred nanometers. They were mostly found at the air-water interface or the carbon support film, indicating that they preferred to bind to distinct surfaces (*i.e.,* hydrophobic surfaces). Along their axis, they showed a very regular stripe pattern with 55 Å distancing, similar to the hairpin array distances found in Type I tubes and carpets (**Suppl. Fig. 8E**). Vipp1 assemblies on larger vesicles (>200 nm) are mostly present in either small patches inducing bulges or tubes/rings creating membrane protrusions with positive membrane curvature (gallery of tomograms in **Fig. 5C**). Intermediate vesicles (around 100 nm) are almost fully covered with Vipp1 carpets, showing the characteristic spike pattern while they often have irregular shapes or large bulges. In some cases, we found intricate membrane networks with very high Vipp1 coverage and regions with high local membrane curvature. At last, we found small vesicles (15 – 80 nm) fully covered with Vipp1. Most small vesicles that were not encapsulated were covered with Vipp1 including the regular spike patterns. Notably, we could not clearly distinguish between very small Vipp1-membrane assemblies (<15 nm) and Vipp1 ring complexes with engulfed membranes in our tomograms.

For a more quantitative analysis of the observed Vipp1 membrane assemblies, we determined the membrane curvature for 123 tomograms (**Suppl. Fig. 9A**). In addition, for each segmented membrane voxel region we checked for Vipp1 voxel presence and for each curvature we computed an occupancy share with zero representing empty and one fully covered membranes (**Fig. 6A** right, **Suppl. Fig. 9B**). Based on the here determined values, membrane curvature correlates with Vipp1 occupancy (**Fig. 6A** left). The membrane occupancy increases linearly with increasing membrane curvedness between curvedness values of 0.02 to 0.06. For high curvature regions, *i.e.,* for curvednesses larger than 0.06, the graph indicated a saturation of Vipp1 membrane binding for high curvature regions. To support the assumption that Vipp1’s affinity is higher for liposomes with high curvature, we compared binding of the protein to liposomes of different sizes (**Fig. 6B, Suppl. Fig. 9C**). Employing tryptophan fluorescence spectroscopy, we observed pronounced changes in the fluorescence emission spectra when the protein was incubated with increasing concentrations of sonified liposomes with an average diameter of ∼55 nm as opposed to extruded liposomes with an average diameter of ∼144 nm where hardly any change in tryptophan fluorescence was detected. This observed difference indicates that Vipp1 preferrably binds to small liposomes with high membrane curvature supporting the apparent Vipp1 coverage dependence on the curvature observed in the tomograms.

**Figure 6:**
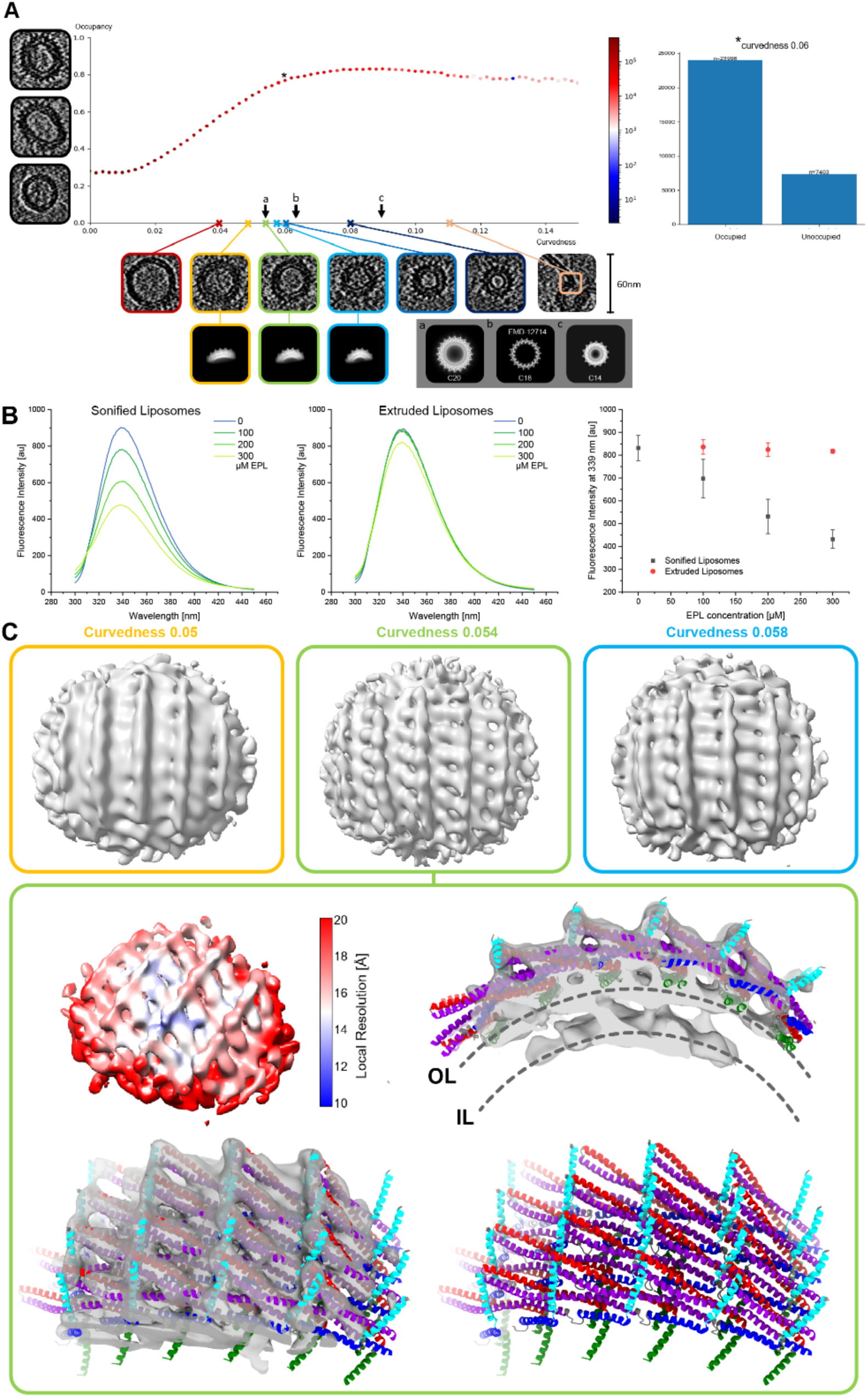
Vipp1 membrane curvature analyses and corresponding subtomogram averages. A: Membrane curvature in relation to occupancy representing the Vipp1 membrane coverage. Left: Example images of different occupancies with fully covered, partly covered, and uncovered vesicles. Bottom: Example images of vesicles and other shapes with specified curvedness. 2D averages of subtomogram averages corresponding to the indicated curvedness. Bottom right: 2D average of C14, C18, and C20 ring as comparison. Right: Exemplary examination for all segmentation points with a curvedness of 0.06. **B:** Tryptophan fluorescence spectroscopy of Vipp1 in the presence of sonified (∼55 nm diameter) or extruded (∼144 nm) liposomes. Solely upon interaction of refolded Vipp1 with sonified liposomes, a pronounced decrease in quantum yield is observed for small liposomes. Error bars represent SEM, n=4. **C:** Subtomogram averages for curvedness 0.05, 0.054, and 0.058 (top). Local resolution map yielding an average resolution of 18 Å (left). Density maps with fitted atomic model based on the C20 symmetrization (adapted from PDB:7O3Z). OL: outer leaflet, IL: inner leaflet.

To further investigate the structure and organization of Vipp1 in the membrane-attached carpets, we excised subvolumes of 50 nm from the tomograms and performed subtomogram averaging of Vipp1-occupied membrane cubes, excluding small versicles (<15 nm) and tubes from the analysis. This way, we were able to determine three structures of Vipp1 carpets for curvedness 0.05, 0.054, and 0.058, at 20, 18, and, 19 Å resolution, respectively (**Fig. 6C, Suppl. Fig. 9D**). The reconstructed structures included density for the inner and outer membrane leaflet in addition to the Vipp1 lattice that was remarkably similar to Vipp1 ring complexes with respect to spike separation distance and monomer stacking distance. To generate an atomic model for our highest resolution structure (curvedness 0.054, 18 Å global resolution), we placed a lattice of a generated C20 monomer assembly (derived from PDB: 7O3Z) guided by the characteristic α1–3 hairpin-to-α5 spike shape in the periphery in the density (**Suppl. Fig. 9E**). In accordance with previously described Vipp1 polymers, Vipp1 monomers form the typical α1–4 contacts in the core and α1–3 hairpin-to-α5 contacts in the periphery. Helix α0 presumably mediates membrane contacts. The observation that it was not possible to determine a structure for lower curvature Vipp1 carpets, despite abundant particle data, may indicate less membrane coverage and regularity in the arrangement of low curvature Vipp1 carpets, in agreement with recent AFM measurements on flat membrane surfaces (Junglas et al., 2020a). Finally, our observations indicate a strong correlation between membrane curvature and the degree of Vipp1 membrane coverage.

## Discussion

In this study, we elucidated the structures of different Vipp1 assemblies bound to membranes from highly ordered symmetrical assemblies to more loosely organized membrane-covering carpets (**Fig. 1**). The best-organized states were helical tubes that occurred in two different architectural types and stacked-ring Vipp1 assemblies of cyclical symmetries, all of which included lipid bilayer density in the lumen (**Fig. 2**, **Fig. 3**). Membrane tubulation of Vipp1 is largely mediated by helix α0, as truncated Vipp1(α1-6) assembled into helical tubes that did not internalize any lipid membrane in the lumen (**Suppl. Fig. 4 and 5**). By generating the plasticity-restrained Vipp1 dL10Ala mutant, we could narrow down the variability of Vipp1 assemblies and solve the structure of two Vipp1 tubes at 3.0 Å resolution. These structures revealed that one positively charged face of helix α0 serves as a central contact point to hold helices α0, α1 and α4 from four adjacent subunits in place while the hydrophobic face can interact with acyl chains of engulfed membrane lipids (**Fig. 4**). Less well-ordered Vipp1 assemblies included membrane-free spiraling ribbons as well as membrane-associated loose coats, patches, and carpets (**Fig. 5**). Membrane coverage of Vipp1 correlates with higher membrane curvature leading to the observed more ordered symmetrical Vipp1 complexes (**Fig. 6**).

Our Vipp1-containing membrane preparations revealed a remarkable diversity of membrane-bound Vipp1 structures suitable for structure determination, which had not been observed previously. Past studies worked with pre-formed ring complexes incubated with membranes (Heidrich et al., 2016; Hennig et al., 2015; Junglas et al., 2020a; Liu et al., 2021; Theis et al., 2019). These studies suggested that preformed *Synechocystis* Vipp1 ring complexes in solution bind to membranes and afterward disassembled into carpet structures, while *Chlamydomonas reinhardtii* Vipp1 formed rod structures with internalized membranes. Clearly, the observations depend on the Vipp1 homolog and the exact preparation conditions used in the different studies. The observation that lipids are often copurified with pre-formed Vipp1 assemblies isolated from *E. coli* after heterologous expression could also lead to a decrease in apparent membrane binding and membrane remodeling (Gupta et al., 2021; Otters et al., 2013). Likely, pre-formed Vipp1 ring complexes are less reactive as they are already present in a polymeric form, as has been observed for the closely related *Syn*PspA (Junglas et al., 2021). Upon interaction with membranes, preformed Vipp1 polymers may just slowly disassemble due to slightly favoring protein-membrane interactions. Indeed, in a recent analysis of assembled *Synechocystis* Vipp1 ring complexes binding to a solid-supported bilayer, disassembly of the ring complexes on the membrane surface was observed via AFM (Junglas et al., 2022, 2020a). Due to the solid support, the membrane may not be as flexible as a not-tethered membrane of liposomes, thus allowing the formation of carpets but not of membrane-attached rings or tubes. Now, we purified Vipp1 under denaturing conditions and refolded it in the presence of lipids. We identified refolding in the presence of membranes as a condition more capable of membrane remodeling giving rise to the observed diversity of Vipp1 membrane assemblies while keeping their native ESCRT-III fold intact.

The here determined Vipp1 stacked-ring assemblies closely resemble previously observed membrane-free ring complexes (Gupta et al., 2021; Liu et al., 2021), except for an internalized continuous membrane tube connecting the rings. The ring complexes are built by six unique layers of different monomer structures. The monomers differ at the hinge 2 and 3 regions between helices α3 and α4, α4 and α5, and the length of α4. The angle of the α1–3 hairpin changes from the bottom to the top layers (see **Fig. 3**). At the bottom layer, the hairpin has a nearly 90° angle to the ring axis, while by approaching the top layer, the angle decreases to ∼60-70°. This effect is more pronounced at smaller ring complex diameters (Gupta et al., 2021; Liu et al., 2021). In contrast to previous studies based on bulk FRET and fluorescence measurements (Heidrich et al., 2018), our stacked-ring assembly structures now clearly show that Vipp1 rings are joined in a polar head-to-tail fashion (see **Fig. 3A**). This head-to-tail interaction is mediated by the docking of the α1–3 hairpin of the lower ring to α5 of the upper ring. This interaction mode is in good agreement with the observation that mutation of conserved residues at the tip of the α1–3 hairpin prevents Vipp1 ring stacking (Saur et al., 2017). Interestingly, it has been shown that the addition of Mg^2+^ can induce ring stacking in the absence of membranes (Heidrich et al., 2018; Saur et al., 2017), suggesting that Mg^2+^ may stabilize the α1–3 hairpin-to-α5 interaction. The membrane engulfed in stacked-ring assemblies exhibits a notable feature as it does not form a uniform tube diameter throughout the stacked-ring assembly. Instead, the membrane follows the tapered wall of the ring complex and is constricted at the sites where two rings are touching and at the “open” ends of the stacked-ring assemblies. Consequently, the membrane undergoes a soft kink of approx. 120° (see **Fig. 3C and D**). The high local curvature renders the membrane suitable for spontaneous fusion with other membranes, in particular at the “open” ends. This way, one ring complex may connect to another membrane-filled ring complex and thereby elongates the stacked-ring assembly or fuse with a free membrane. In fact, the membrane outline inside stacked-ring assemblies closely resembles the typical hour-glass shape of two fusing membranes that are caged in that highly reactive state by the Vipp1 lattice. The observed membrane curvature changes within the structures are formed by the protein cage and could also result in the local accumulation of a specific lipid subpopulation and redistribution of the used EPL mixtures. Together, our data support the concept that the protein assembly structure stabilizes possible membrane transition states highly suitable for Vipp1-mediated membrane fusion, involving the induction of high local membrane curvature within the Vipp1 rings (Hennig et al., 2015; Liu et al., 2021; Saur et al., 2017; Siebenaller et al., 2019). The here determined two types of helical rods represent unique Vipp1 structures not described before. In contrast to the Vipp1 rings, the monomer orientation of the α1–3 hairpin with respect to the tube axis is the same over the whole length of the tubes. Nevertheless, the monomer orientation among the tubes differs: in the Type I tubes, the α1–3 hairpins are oriented parallel to the tube axis, while in both Type II tubes, these are tilted approximately 45° relative to the tube axis. Within the Vipp1 ring complexes, the α1–3 hairpin orientation changes gradually from approx. 90° at the bottom to approx. 60-70° at the top. Remarkably, from rings over Type II tubes to Type I tubes the α1–3 hairpins accommodate a wide range of orientations along the tubulated membrane axis from perpendicular to parallel (from 90° to 0°) while essentially maintaining the same intermolecular contacts to stabilize the assembly (**Fig. 7A**). Interestingly, in our plasticity-restrained mutant, the structural diversity of membrane interacting assemblies was limited to predominantly Type II tubes, *i.e.*, to hairpins aligned 45° relative to the tube axis. Therefore, it can be expected that the intrinsic structural plasticity and conformational flexibility of WT Vipp1 is required to sample different conformations and transition states that assist in membrane remodeling. This kind of structural orientation change appears to be a unique feature of Vipp1, as the other ESCRT-III proteins with known structures on membrane tubes (*i.e., Syn*PspA, CHMP2A+CHMP3, CHMP1B, and CHMP1B+IST1) have a fixed orientation relative to the membrane axis (Azad et al., 2023; Junglas et al., 2021; Nguyen et al., 2020) (**Suppl. Fig. 9F**).

**Figure 7:**
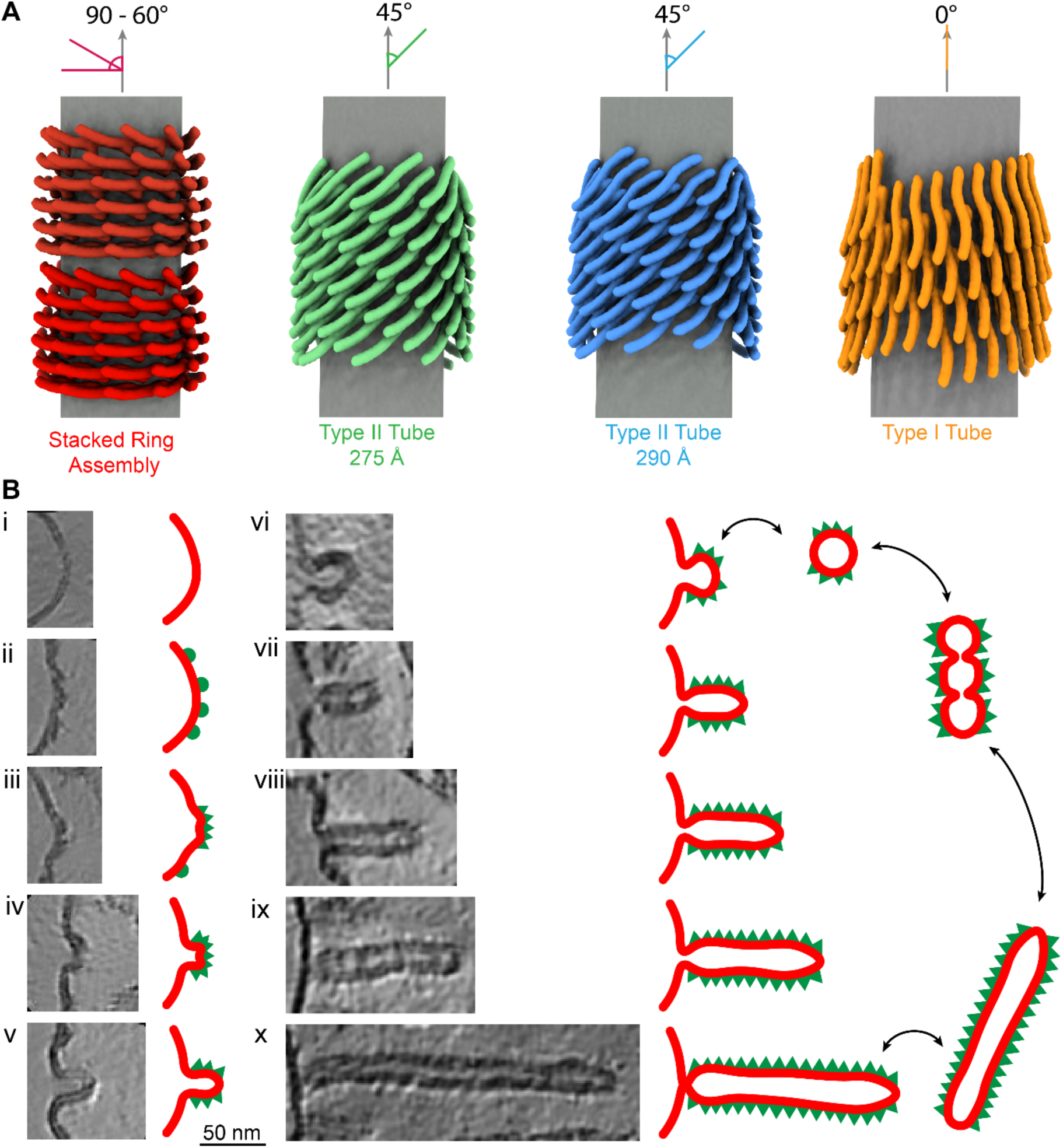
Orientation of the Vipp1 hairpin and sorted tomographic slices of Vipp1 membrane interactions. A: Schematic view of Vipp1 *α*1-3 hairpins relative to the membrane axis in stacked-ring assembly, Type II, and Type I tubes. **B:** Based on the identified structures, we sorted them with regard to Vipp1 membrane interactions and the degree of membrane deformation. Initially, Vipp1 binds to free membranes (*i*). When the Vipp1 monomers reach a critical local concentration on the membrane surface (*ii*), they start to oligomerize and form a membrane bulge (*iii*). Vipp1 induces membrane curvature upon oligomerization (*iv*). The induced curvature can result in the formation of a budding Vipp1 oligomer with enclosed membrane (*v*). This assembly can either bud off the membrane to produce rings (*vi*) or grows further to extended rod structures (*vii – x*). The rods can reside at the host membrane or bud off to produce Vipp1-coated membrane tubes. The budded rings can fuse to form stacked rings that finally relax to Vipp1-coated membrane tubes. Please note that this model is limited by the observation of Vipp1 assemblies in micrographs and sorting them into logical sequence.

The determined structures of stacked-ring assemblies and helical tubes revealed an internalized lipid bilayer. The bilayer thickness in all three Vipp1 tubes remained nearly identical at 35 Å, although the inner leaflet radius changed with the tube diameter. This observation differs from the previous reports about CHMP1B and CHMP1B+IST1 (co)polymers where the bilayer thickness decreased with decreasing tube diameter (Nguyen et al., 2020). The bilayer thickness in the Vipp1 stacked-ring assemblies, however, changed only slightly with the increasing diameter of the rings. This observation may be a result of the high local membrane curvature within the stacked-ring assemblies. Similar to other ESCRT-III proteins, the membrane contact in the Vipp1 tubes and rings is mediated either by an additional α-helix (*i.e.,* α0) in the case of Vipp1 and *Syn*PspA or a short unstructured extension of helix α1 in the case of CHM1B or CHMP2A+CHMP3 (Azad et al., 2023; Junglas et al., 2021; Moss et al., 2023; Nguyen et al., 2020). However, in contrast to *Syn*PspA helix α0 that protrudes towards the bilayer (Junglas et al., 2021), Vipp1 helix α0 is lying flat in the membrane plane and appears to be partially submerged into the outer bilayer leaflet. The 3.0 Å resolution structure of a plasticity-restrained Vipp1-dL10Ala mutant revealed the amphipathic positioning of helix α0 with the hydrophobic face directed into the tube lumen to potentially interact with hydrophobic acyl chains while the positive helix face interacts with the inner Vipp1 ring/tube surface and may be capable of interacting with lipid head groups.

Interestingly, truncation of helix α0 prevents the internalization of membranes into the lumen of the assemblies and/or formation of assemblies on the membrane. This observation is in line with the reports that the isolated helix α0 of Vipp1 binds to membranes (McDonald et al., 2017, 2015). Although it has been reported that truncated versions of Vipp1 lacking helix α0 are still capable of membrane binding (Thurotte and Schneider, 2019), our micrographs and cryo-EM maps show that Vipp1(α1-6) does not engulf membranes in the rod lumen and does not assemble on membranes while it may still be capable of binding lipids in other conditions. Furthermore, mutation of the conserved residues F4 and V11 in the hydrophobic face of helix α0 led to severely decreased high-light resistance and TM swelling of *Synechocystis* cells (Gupta et al., 2021), possibly because Vipp1 could not properly interact with membranes anymore. Thus, we conclude that helix α0 is the membrane anchor of Vipp1, in line with the suggested N-terminal ANCHR motif of some eukaryotic ESCRT-III proteins (Snf7, CHMP2B, CHMP3, CHMP2A, CHMP1B) (Azad et al., 2023; Bodon et al., 2011; Buchkovich et al., 2013; Moss et al., 2023; Nguyen et al., 2020).

Moreover, the truncation of helix α0 also affects the assembly state of Vipp1. Vipp1 prefers to form rods over rings when residues conserved in helix α0 are mutated (Gupta et al., 2021), or exclusively forms rods, when helix α0 is completely absent (this study and (Thurotte and Schneider, 2019)). Additionally, the monomer orientation and rod symmetry differ from the rods observed upon incubation of the full-length protein with membranes while the Vipp1(α1-6) rods adopt a *Syn*PspA rod-like symmetry and orientation and cover a large range of diameters (Junglas et al., 2021). It has been suggested that columns of helix α0 in the lumen of Vipp1 rings stabilize the ring structure versus rod formation (Gupta et al., 2021). Nevertheless, the same or similar intermolecular interactions of helix α0 with another α0 and α1+4, as observed previously in Vipp1 rings (Gupta et al., 2021; Liu et al., 2021), are present in Type I and II tubes. In the PspA-like Vipp1(α1-6) rods, however, a register shift and a larger distance to the next layer of monomers would make it impossible for helix α0 to reach α1+4 of the monomer in the adjacent layer. Thus, the presence and absence of helix α0 coordinates different assembly states of Vipp1, and interactions of helix α0 stabilize particular polymeric Vipp1 structures. Our 3.0 Å Vipp1 tube structure now reveals that D5, R6, R9, R12, and E23 of helix α0 interact with three different subunits via α4 (R170), α4 (K174) and α1 (D35) to hold these helices in place, thereby aligning the interacting subunits/helices in an orientation favoring rings and Type I/II tubes over the PspA-like assembly, in addition to described lipid interaction properties.

In addition to the well-ordered symmetrical structures, we were able to determine the structure of membrane-bound Vipp1 carpets on the molecular level. The determined low-resolution Vipp1 carpet structures were reminiscent of wider Vipp1 ring complexes, presumably with an approx. 90° rotation of the hairpin relative to the membrane axis. Formation of Vipp1 carpets is supposed to play an important role in Vipp1 membrane stabilization/protection and has been observed on flat solid-supported bilayers *in vitro* and presumably on TMs *in vivo* (Gutu et al., 2018; Junglas et al., 2020a; Junglas and Schneider, 2018). Additionally, *in vivo* observations of Vipp1 rods/tubes and (presumably) carpets, but not ring complexes, suggest that these are equally physiologically relevant or stable forms of Vipp1 polymers (Gupta et al., 2021; Gutu et al., 2018; Junglas and Schneider, 2018). We identified Vipp1 carpet assemblies on low curvature as well as highly curved membranes while we found higher Vipp1 coverage on high-curvature membranes, in line with the observation that Vipp1 prefers binding to highly curved membranes (see **Fig. 6**). Strikingly, our subtomogram averages of the carpet structures suggest a monomer arrangement similar to the one found in Vipp1 rings, except that the assemblies do not close into a ring. Instead, the lattice is relaxed to adapt to lower membrane curvatures and extended to cover larger areas. The abundance of membrane-bound Vipp1 structures (loose coats, patches, carpets, and rings) increased with increasing membrane curvature and reached a plateau at a curvature corresponding to an approximate vesicle diameter of 30 nm. Higher-diameter (lower curvature) membranes are less frequently occupied with Vipp1 assemblies, in line with the observation that membrane-bound Vipp1 is concentrated at high curvature regions of the TM *in vivo* (Gutu et al., 2018). Together, we observed that the local Vipp1 molecule concentration on the membrane affects the assembly state and thereby critically modulates membrane curvature (and *vice versa*).

In light of our findings, we propose the following sequence for Vipp1 membrane interactions (**Fig. 7B**): (*i*) Vipp1 monomers/small oligomers bind to and (*ii*) accumulate on low-curvature membranes in loose coats until a critical concentration is reached locally. (*iii*) At these membrane areas, Vipp1 monomers oligomerize in the plane of the membrane and the formed assembly patches start to induce curvature on the membrane, ultimately resembling the determined carpet structures. As Vipp1 preferentially binds to curved membranes, *i.e.,* membrane binding induces curvature, (*iv, v*) Vipp1 bulges start to grow on the membrane increasing the local curvature, stacking of oligomers away from the membrane and thereby internalizing the membrane through helix α0 interactions. (*vi*) This way, the initial bulges further emerge into Vipp1 ring and (*vii – x*) stacked-ring complexes and subsequently elongate into rods that finally bud off the membrane. The transition of the emerging Vipp1 assembly is accompanied by a changed orientation of the Vipp1 hairpin α1-3 from 90° at the low curvature carpet to approx. 0° at the highest curvature with respect to the emerging tube axis. The hour-glass shape of the stacked-ring assemblies will promote the transition from a flat membrane geometry to a highly curved membrane. When the Vipp1 assemblies constrict the internalized membrane beyond a certain threshold, the host membrane is pinched off and the assemblies can leave the membrane surface. Additionally, rings and short rods that budded off the membrane still contain highly curved membranes at their ends, prone to fuse with available free membranes to spawn new Vipp1 assembly sites, or with other ring complexes to form stacked rings. The driving forces for this process are determined by the tendency of Vipp1 to form highly ordered tubular lattices upon membrane interaction and thereby stabilizing curved membranes. As our data lack any temporal resolution, we cannot exclude that the process is running in the other direction or back and forth in both directions depending on the exact experimental conditions. In fact, previous studies suggested that Vipp1 ring complexes in solution bind to membranes, subsequently disassemble (Heidrich et al., 2016), and transform into carpet structures (Junglas et al., 2020a). Moreover, time-resolved structural research will be required to delineate the temporal order and other short-lived intermediates to comprehensively describe the structural states involved in this Vipp1-mediated membrane remodeling process.

## Material and Methods

### Expression and purification of Vipp1

Full-length Vipp1 (*sll0617*) of *Synechocystis* sp. PCC 6803 and associated mutants (Vipp1(α1-6) (truncation of helix α0 (aa 1-23)), Vipp1 dL10 (truncation of aa 157-166), and Vipp1 dL10Ala (mutation of aa 157-166 to Ala)) were heterologously expressed in *E. coli* C41 cells in TB medium using a pET50(b) derived plasmid with a C-terminal His-tag and 3C-protease cleavage site. For purification of Vipp1 and Vipp1(α1-6) under denaturing conditions, cells were resuspended after protein expression in lysis buffer containing 6 M urea (10 mM Tris-HCl pH 8.0, 300 mM NaCl) supplemented with a protease inhibitor. Cells were lysed in a cell disruptor (TS Constant Cell disruption systems 1.1 KW; Constant Systems). The crude lysate was supplemented with 0.1% (*v*/*v*) Triton X-100 and incubated for 30 min at room temperature. Subsequently, the lysate was cleared by centrifugation for 15 min at 40,000 g. The supernatant was applied to Ni-NTA agarose beads. The Ni-NTA matrix was washed with lysis buffer and lysis buffer with additional 10 mM imidazole. The protein was eluted from the Ni-NTA with elution buffer (10 mM Tris-HCl pH 8.0, 1000 mM imidazole, 6 M urea). The fractions containing protein were pooled, concentrated (Amicon Ultra-15 centrifugal filter 10 kDa MWCO), and stored at -20 °C. For cleavage of the C-terminal His-tag, the protein was diluted 1:1 in cleavage buffer (10 mM Tris-HCl, 300 mM NaCl, 1mM DTT, 0.1 mM EDTA) and dialyzed overnight against cleavage buffer together with 3C-protease, including three buffer exchanges. After the cleavage reaction, the protein was diluted 1:3 with 10 mM Tris-HCl pH 8.0, 8 M urea, 200 mM NaCl, and 7 mM imidazole and applied to the Ni-NTA matrix. The Ni-NTA matrix was washed with 10 mM Tris-HCl pH 8.0, 6 M Urea, 100 mM NaCl, and 4 mM imidazole. Flow through and wash fractions containing the cleaved protein were pooled, concentrated (Amicon Ultra-15 centrifugal filter 10 kDa MWCO), and stored at -20 °C. Protein concentrations were determined by measuring the absorbance at 280 nm of Vipp1 diluted in 4 M guanidine hydrochloride using the respective molar extinction coefficient calculated by ProtParam (Gasteiger et al., 2005).

### Liposome preparation and lipid reconstitution

Chloroform dissolved *E. coli* polar lipid (EPL) extract was purchased from Avanti polar lipids (*Birmingham, AL, USA*). Lipid films were produced by evaporating the solvent under a gentle stream of nitrogen and vacuum desiccation overnight. For Vipp1 reconstitution with lipids, urea-unfolded Vipp1 was added to the EPL film (see **Suppl. Table 1**) and incubated with shaking for 20 - 30 min at 37 °C until the lipid film was resolved. Subsequently, the mixture was dialyzed overnight against 10 mM Tris-HCl pH 8.0 (8 °C, 20 kDa MWCO) including three buffer exchanges to refold the protein.

For analysing membrane binding of Vipp1 helix α0 peptides, buffer was added to the EPL film to a final lipid concentration of 8 mM and incubated at 37 °C with shaking for 30 min. The liposomes were then prepared by sonication (Branson Tip sonifier).

### Characterization of the refolded protein

Vipp1 prepared as described above was refolded by dialysis against 10 mM Tris-HCl pH 8.0 over night, with three buffer exchanges. Protein concentrations were determined based on the protein absorption at 280 nm as described above. For comparison, Vipp1 purified under native conditions (*e.g.* (Hennig et al., 2015)) was studied. The protein concentration was adjusted to match the concentrations of the refolded protein based on a comparison of the tryptophan fluorescence intensity at 340 nm determined in 6 M urea. The thermal stability was studied by monitoring the change in secondary structure through circular dichroism (CD) spectroscopy (J-1500, Jasco, Pfungstadt, Germany). The CD spectrum between 210 and 230 nm was measured at different temperatures (2°C temperature steps, 1 nm wavelength steps, overall mean temperature ramp 10°C/h). Samples of each protein were prepared in duplicate and measured in parallel in a 6-cell holder. Interaction of the refolded protein with preformed liposomes was monitored by following changes in tryptophan fluorescence. To this end, an EPL film, prepared as described above, was rehydrated with Tris-buffer. Unilamellar liposomes were either formed by sonication (Branson Tip sonifier), to yield small liposomes or by extrusion through a 200 nm filter (Avanti Polar Lipids, Birmingham, AL, USA). The size of the liposomes was determined by dynamic light scattering (Zetasizer Pro, Malvern, Kassel, Germany). Based on the z-average, the extruded liposomes had a diameter of 143 ± 0.5 nm, and the sonified liposomes 56 ±0.44 nm. A concentration of 3 µM protein was incubated with liposomes for 45 min at room temperature. Fluorescence emission was determined from 300 nm to 450 nm at 25°C after excitation at 280 nm, with both excitation and emission slit widths set to 2.5 nm (FP-8500, Jasco, Pfungstadt, Germany).

### Membrane binding of Vipp1 helix α0 peptides

The peptides Vipp1 helix α0 wt [MGLFDRLGRVVRANLNDLVSKAED], Vipp1 helix α0 R6A_R9A [MGLFDALGAVVRANLNDLVSKAED], Vipp1 helix α0 F4A_V11A

[MGLADRLGRVARANLNDLVSKAED] and Vipp1 helix α0 amph(-) [MGAADRAGRAARANANDAASKAED] were purchased from PSL GmbH (Heidelberg, GER). The changes in secondary structure upon membrane binding were determined by circular dichroism (CD) spectroscopy (J-1500, Jasco, Pfungstadt, Germany). 40µM peptide (10 mM Tris, pH 8.0) was incubated with increasing amounts of liposomes (up to 4 mM EPL). After incubation at room temperature for 15 min, spectra were recorded between 190 and 250 nm in 1 nm steps at 25°C. Three individual samples were measured for each sample, with 12 accumulations per sample. The mean and standard deviation of the value from the three samples at each lipid concentration were used to create a binding curve..

### Negative staining electron microscopy

For negative-staining electron microscopy, 3 µL of the sample was applied to glow-discharged (PELCO easiGlow Glow Discharger, Ted Pella Inc.) continuous carbon grids (CF-300 Cu, Electron Microscopy Sciences). The sample was incubated on the grid for 1 min. Then, the grid was side-blotted using filter paper, washed with 3 µL water, stained with 3 µL 2% uranyl acetate for 30 s, and air-dried. The grids were imaged with a 120 kV Talos L120C electron microscope (ThermoFisher Scientific/FEI) equipped with a CETA camera at a pixel size of 2.49 Å/pixel (57 kx magnification) at a nominal defocus of 1.0 to 2.5 µm.

### Electron cryo-microscopy

Grids were prepared by applying 4 μL sample (with or without 5 nm gold fiducials) (**Suppl. Table 1**) to glow-discharged (PELCO easiGlow Glow Discharger, Ted Pella Inc.) Quantifoil grids (R1.2/1.3 Cu 200 mesh, Electron Microscopy Sciences). The grids were plunge-frozen in liquid ethane using a Leica EM GP2 plunge freezer set to 80% humidity at 10 °C (sensor-guided back-side blotting, blotting time 3 to 5 s). Movies were recorded in under-focus on a 200 kV Talos Arctica G2 (ThermoFisher Scientific) electron microscope equipped with a Bioquantum K3 (Gatan) detector or a 300 kV Titan Krios G4 (ThermoFisher Scientific) electron microscope equipped with a Biocontinuum K3 (Gatan) and a Falcon4i (ThermoFisher Scientific) detector operated by EPU (ThermoFisher Scientific) or Tomo (ThermoFisher Scientific). Tilt series for cryo-ET were collected at -60° to 60° with 3° in a dose-symmetric scheme. Tilt images were acquired at a magnification of 53 kx (pixel size 1.7 Å) with a nominal underfocus of 3.0 µm. The total dose for each tilt series was 152 e^−^/Å^2^. A total of 176 tilt series were collected (summarized in **Table 1** and **Suppl. Table 1** for data acquisition details of the single-particle and cryo-electron tomography acquisitions).

### Single-particle image processing and helical reconstruction

Movie frames were gain-corrected, dose-weighted, and aligned using cryoSPARC Live (Punjani et al., 2017). Initial 2D classes were produced using the auto picker implemented in cryoSPARC Live. The following image processing steps were performed using cryoSPARC. The classes with most visible detail were used as templates for the filament tracer. For the Vipp1 EPL dataset, the resulting filament segments were extracted with 600 px box size (Fourier cropped to 200 px) and subjected to multiple rounds of 2D classification. The remaining segments were sorted by filament class: (*i*) stacked rings, (*ii*) Type I tubes, and (*iii*) Type II tubes. For each filament class, the remaining segments were reextracted with a box size of 400 px (Fourier cropped to 200 px) and subjected to an additional round of 2D classification. The resulting 2D class averages were used to determine filament diameters and initial symmetry guesses in PyHI (Zhang, 2022). Initial symmetry estimates were validated by helical refinement in cryoSPARC and selection of the helical symmetry parameters yielding reconstructions with typical Vipp1 features and the best resolution. Subsequently, all segments for each filament class were classified by heterogeneous refinement and followed by 3D classifications using the initial helical reconstructions as templates. The resulting helical reconstructions were subjected to multiple rounds of helical refinement including the symmetry search option. Reconstructions of the stacked rings were treated similarly, but instead of helical symmetry, the respective rotational symmetry was applied in helical reconstruction jobs (C11-C14). Segments with poor visible details were discarded by heterogeneous refinement. For the final polishing, the segments were re-extracted at 600 px with Fourier cropping to 400 px (stacked rings and Type I tubes) or at 500 px without Fourier cropping (Type II tubes). The Vipp1(α1-6) and Vipp1(α1-6) EPL datasets were preprocessed as described above. The filament segments were extracted with 700 px box size (Fourier cropped to 200 px) and subjected to multiple rounds of 2D classification. The following image processing steps were identical to the above-described workflow of helical reconstruction of Type I and Type II filaments. Bad segments were discarded by heterogeneous refinement. For the final polishing, the segments were reextracted at 700 px with Fourier cropping to 350 px.

The Vipp1 dL10Ala EPL dataset was preprocessed using WARP (Tegunov and Cramer, 2019) (motion correction, binning to physical pixel size, and CTF estimation) and further processed in cryoSPARC as described above. The filament segments were extracted with 500 px box size (Fourier cropped to 200 px) and subjected to multiple rounds of 2D classification. The following image processing steps were identical to the above-described workflow of helical reconstruction of Type I and Type II filaments. Poor segments were discarded by heterogeneous refinement or 3D classification. For the final polishing, the segments were reextracted at 500 px without Fourier cropping. The ring structures were picked using a template picker and extracted at 600 px with Fourier cropping to 200 px. The extracted particle stack was cleaned after multiple rounds of 2D classification. Rings with different rotational symmetries were sorted by multi-class *ab-initio* reconstruction and further classified by multiple rounds of heterogenous refinement and *ab-initio* reconstruction with imposed rotational symmetry (C15-C20). Poor particles were discarded by heterogeneous refinement. For the final polishing, the particles were reextracted at 600 px with Fourier cropping to 450 px and subjected to nonuniform refinement with imposed symmetry (C15-C20). The local resolution distribution and local filtering for all maps were performed using cryoSPARC (as shown in **Suppl. Fig. 2A, 3A, 4E, 6D, and 7A**). The resolution of the final reconstructions was determined by Fourier shell correlation (auto-masked, FSC=0.143).

### Cryo-electron tomography image processing

Tilt image frames were gain-corrected, dose-weighted, and aligned using WARP (Tegunov and Cramer, 2019). The resulting tilt series were aligned, 8x binned, and reconstructed by the weighted back projection method using AreTomo (Zheng et al., 2022). For segmentation, the reconstructed tomograms were filtered with a recursive exponential filter followed by a non-local means filter using Amira (ThermoFisher Scientific). The filtered tomograms were segmented in Dragonfly (Object Research Systems) by progressively training a U-Net with an increasing number of manually segmented tomogram frames (5-15). Then the trained U-Net was used to predict those features of the tomogram. For visualization, the segmentation was cleaned up by removing isolated voxels of each label group (islands of <150 unconnected voxels). The resulting segmentation was imported to ChimeraX (Pettersen et al., 2021) for 3D rendering. For quantitative analysis of membrane curvature, 8 tomograms were used in the same manner to train the U-Net, which was then used to predict 123 tomograms included in the analysis. Once the resulting segmentations were coarsely corrected for errors, they were exported as .tif files and assembled into .mrc files for further analysis. For the membrane segmentations, the surface morphometrics toolkit (Barad et al., 2023) was used to determine curvature and membrane normals. The resulting coordinates and their corresponding curvature estimates were used to filter for adjacent carpet segmentations, accepting a distance up to 50 Å. To minimize errors, coordinates close to tomogram edges were excluded. For further analysis, coordinates were sorted into fixed-size bins according to their curvedness and for each bin, the ratio of membrane coordinates with a present proximal carpet segmentation was determined resulting in the corresponding occupancy value between 0 and 1 (0 no coverage, 1 full coverage).

For subtomogram averaging, Relion4 (Zivanov et al., 2022) was used. CTF estimation was conducted with ctffind4 (Rohou and Grigorieff, 2015). As initial particles, the coordinates resulting from the previously described membrane alaysis were used. Coordinates were selected for the presence of Vipp1 carpet structures (within a radius of 100 Å) and a minimal inter-particle distance of 15 Å was enforced to avoid excessive overlaps. The orientation of particles was initially estimated using their corresponding membrane normal to restrain angular searches for all subsequent refinement procedures. To create an initial model from a subset of homogenous particle picks, all coordinates were clustered according to their corresponding membrane curvature, using the Scikit-Learn implementation of k-means. After particle averaging and several rounds of 3D classifications and refinements in 4x binning, a low-resolution model emerged. This model was then used as an initial reference and from here on, all particles were included. Multiple rounds of 3D classifications and refinements for cleaning up the dataset followed, initially using 4x binning and later on 2x binning. As a further means to exclude poor particles, particles with diverging orientations compared to the majority of particles in close proximity were excluded. Finally, the resulting maps were used to improve predetermined parameters, using Relion’s frame alignment and CTF refinement procedures followed by a final round of refinements. For the highest resolution map, the average curvedness of all points from which the final average was calculated (∼0.054) indicated a substantially higher diameter than the currently existing highest diameter Vipp1 ring model (C18, PDB:7O3Z). Therefore, 7O3Z was used to initially set the distance to the symmetry axis and rotation. To find the correct symmetry parameters (C-symmetry, distance, and rotation in relation to the symmetry axis), the map was fitted in ChimeraX (Pettersen et al., 2021) along the x-axis. The correlation of the original map with versions of itself with different symmetry parameters applied was then calculated, and the correlation was aimed to be maximized. First, translational searches along the x-axis in 1 Å increments were conducted for C18-C22 symmetries. The best correlation for the 0.054 curvedness reconstruction was identified for a C20 symmetry. When optimal distance and symmetry parameters were found, rotational searches were conducted (illustrated in **Suppl Fig. 6E**). Finally the symmetry, shift, and rotational parameters were applied to the reconstruction, and PDB:7O3Z was rigid-body fitted into the resulting map.

### Cryo-EM map interpretation and model building

The handedness of the maps was determined by rigid-body fitting Vipp1 reference structures (PDB:7O3W) into the final maps using ChimeraX (Goddard et al., 2018; Pettersen et al., 2021) and flipped accordingly. For models of the Vipp1 Type I, Type II, and Vipp1 dL10Ala Type IIb tubes, one of the monomers of PDB:7O3W was flexibly MDFF fitted to the 3D reconstructions using ISOLDE (Croll, 2018). The models were subjected to auto-refinement with *phenix.real_space_refine* (Afonine et al., 2018b). The auto-refined models were checked/adjusted manually in Coot (Emsley et al., 2010) and ISOLDE (Croll, 2018) before a final cycle of auto-refinement with *phenix.real_space_refine* (Afonine et al., 2018b). After the final inspection, the models were validated in *phenix.validation_cryoem (*Afonine et al., 2018a*)/Molprobity (*Williams *et al., 2018).* Then the respective helical symmetry was applied to all models to create assemblies of 60 monomers each to generate *biomt* matrices for deposition. For models of the stacked C11 and C12 rings PDB:6ZVR and PDB:6ZVS were used as references (Liu et al., 2021). Side chains were adjusted to the sequence of *Synechocystis sp.* PCC6803 Vipp1 and the resulting models were rigid-body fitted to the maps. For models of the stacked C13 and C14 rings PDB: 7O3W was used as reference (Gupta et al., 2021). The nucleotide was deleted and the monomers were first rigid-body fitted and then flexibly MDFF fitted to the 3D reconstructions using ISOLDE (Croll, 2018). For models of the Vipp1(α1-6) rods PDB: 7O3W was used as a reference (Gupta et al., 2021). Only the central monomer was selected and aa 1-22 were removed. The truncated model was rigid-body fitted to the maps. For each map, the monomers were then flexibly MDFF fitted using ISOLDE (Croll, 2018). The models were subjected to auto-refinement with *phenix.real_space_refine* (Afonine et al., 2018b). The auto-refined models were checked/adjusted manually in Coot (Emsley et al., 2010) and ISOLDE (Croll, 2018) and subjected to a final cycle of auto-refinement with *phenix.real_space_refine* (Afonine et al., 2018b). After the final inspection, the models were validated in *phenix.validation_cryoem (Afonine et al., 2018a)/Molprobity (Williams et al., 2018).* Then the respective helical symmetry was applied to all models to create assemblies of 60 monomers each to generate *biomt* matrices for deposition. The Vipp1 dL10Ala C15 to C20 ring complex models were generated using Vipp1 WT C15 and C16 ring models as reference (PDB: 7O3X and 7O3Y) (Gupta et al., 2021). The nucleotide was removed and the monomers were first rigid-body fitted and then flexibly MDFF fitted to the 3D reconstructions using ISOLDE (Croll, 2018). Image processing, helical reconstruction, and model building were completed using SBGrid-supported applications (Morin et al., 2013). In this manner, from a total of 37 cryo-EM maps determined in 4 samples (3x Vipp1+EPL rods, 4x Vipp1+EPL stacked rings, 10x Vipp1(α1-6) rods, 10x Vipp1(α1-6)+EPL rods, 4x Vipp1 dL10Ala EPL tubes, 6x Vipp1 dL10Ala rings), a total of 17 unique PDB coordinates were refined and deposited originating from 4 samples (listed in **Table 1**). The 4 Vipp1+EPL stacked ring maps and the 6 Vipp1 dL10Ala ring complex maps were not of sufficient quality to refine models. The remaining complementary Vipp1(α1-6) (+EPL) maps were of identical symmetry consisting of either similar or poorer resolution densities and, therefore, atomic model refinement was not pursued further. Diameters of the Vipp1 rod/tube reconstructions were measured as radial density profiles using ImageJ (Rueden et al., 2017). Outer and inner leaflet radii of engulfed membrane tubes were determined from the same radial density profiles at the peak maxima from each bilayer leaflet.

### Quantification and statistical analysis

Data and statistical analysis were performed using OriginPro 2022b (OriginLab Corp., Northampton, USA). Detailed descriptions of quantifications and statistical analyses (exact values of *n*, dispersion and precision measures used, and statistical tests used) can be found in the respective Figures, Figure legends, and Methods section.

## Author contributions

B.J., D.S., and C.S. designed the research. I.R. cloned, expressed, and purified the proteins. B.J. prepared cryo-EM samples and operated the electron microscopes. B.J. and C.S. determined the cryo-EM structures. B.J. built the refined atomic models. B.J. and D.K. reconstructed and segmented the tomograms. D.K. determined the STA structures. B.J., D.K., and C.S. interpreted the protein structures. M.K. analyzed the peptides. N.H. analyzed the isolated proteins. M.K., N.H., and D.S. interpreted the membrane binding data. B.J., D.K., D.S., and C.S. prepared the manuscript with input from all authors.

## Materials availability

All unique and stable reagents generated in this study are available from the Lead Contact with a completed Material Transfer Agreement.

## Data and code availability

The EMDB accession numbers for cryo-EM maps are EMD IDs:

18384, 18421, 18420, 18422, 18423, 18424, 18425, 18426, 18427, 18428, 18429, 18430, 18431, 18432, 18433, 18434, 18435, 18620, 19863, 19864, 19865, 19866, 19899, 19900, 19901, 19902, 19903, 19904

The PDB accession codes for Vipp1 models are PDB IDs:

8QFV, 8QHW, 8QHV, 8QHX, 8QHY, 8QHZ, 8QI0, 8QI1, 8QI2, 8QI3, 8QI4, 8QI5, 8QI6, 9EOM, 9EON, 9EOO, 9EOP

## Acknowledgments

This study was funded by the Deutsche Forschungsgemeinschaft (DFG) (SA 1882/6-1, SCHN 690/16-1) and CRC1551 ‘Polymer concepts in cellular function’ (DFG project number 464588647). The authors gratefully acknowledge the electron microscopy training, imaging, and access time granted by the life science EM facility of the Ernst-Ruska Centre at Forschungszentrum Jülich. The authors gratefully acknowledge the computing time granted by the JARA Vergabegremium and provided on the JARA Partition part of the supercomputer JURECA at Forschungszentrum Jülich (Thörnig, 2021).

## Supplementary Information

**Suppl. Table 1.** Sample Details.

| | *Vipp1*<br>*EPL* | *Vipp1*<br>($\alpha 1-6$) | *Vipp1*<br>($\alpha 1-6$)<br>*EPL* | *Vipp1*<br>(*dL10Ala*)<br>*EPL* |
| --- | --- | --- | --- | --- |
| Protein Conc. | 4.5<br>mg/mL | 2 mg/mL | 2 mg/mL | 4.5 mg/mL |
| Lipid Conc. | 5 mg/mL | - | 5 mg/mL | 5 mg/mL |
| Magnification | 49 kx | 59 kx | 47 kx | 63 kx |
| Pixel size | 1.737 Å | 1.4 Å | 1.7 Å | 1.36 |
| Frames | 40 | 15 | 25 | 30 |
| Total dose | 48 e<sup>-</sup>/Å<sup>2</sup> | 15 e<sup>-</sup>/Å<sup>2</sup> | 25 e<sup>-</sup>/Å<sup>2</sup> | 60 e<sup>-</sup>/Å<sup>2</sup> |
| Defocus range | 1 to 3 μm | 1 to 2 μm | 2 to 3.5<br>μm | 1.25 to 2.75<br>μm |
| Movies | 3,217 | 5,901 | 2,463 | 29,548 |

**Suppl. Figure 1:**
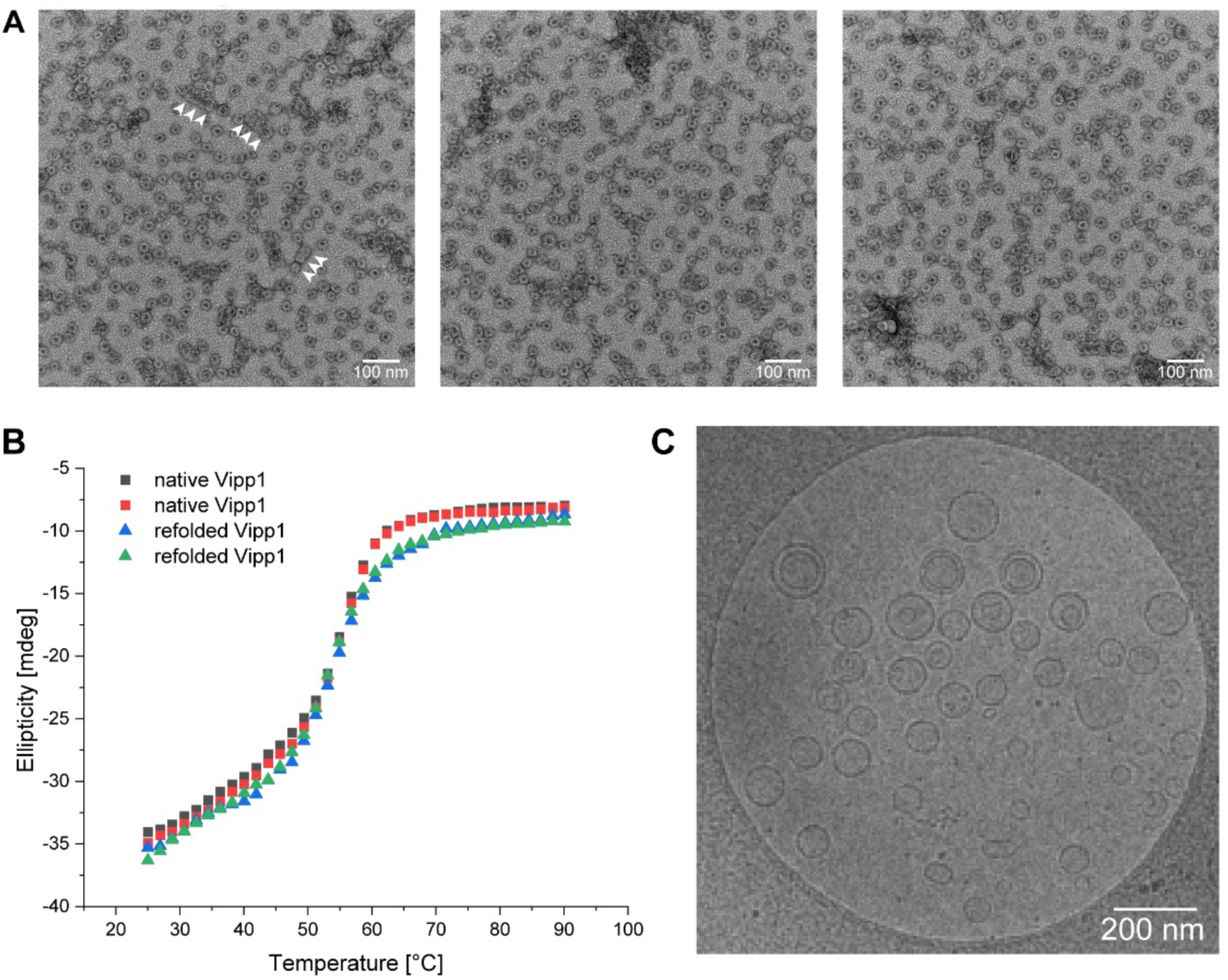
Negative staining EM of Vipp1 preparations. A: Negative staining EM micrographs of Vipp1 in the absence of membranes. In the absence of membranes, Vipp1 forms small oligomers, ring complexes, and stacked-ring assemblies (white arrowheads). **B:** Thermal stability of Vipp1 proteins. Vipp1 with His-tag purified under native conditions (black, red) or Vipp1 purified in the presence of 6 M urea (green, blue) was exposed to increasing temperatures. **C:** Cryo-EM micrograph of EPL liposomes without Vipp1.

**Suppl. Figure 2:**
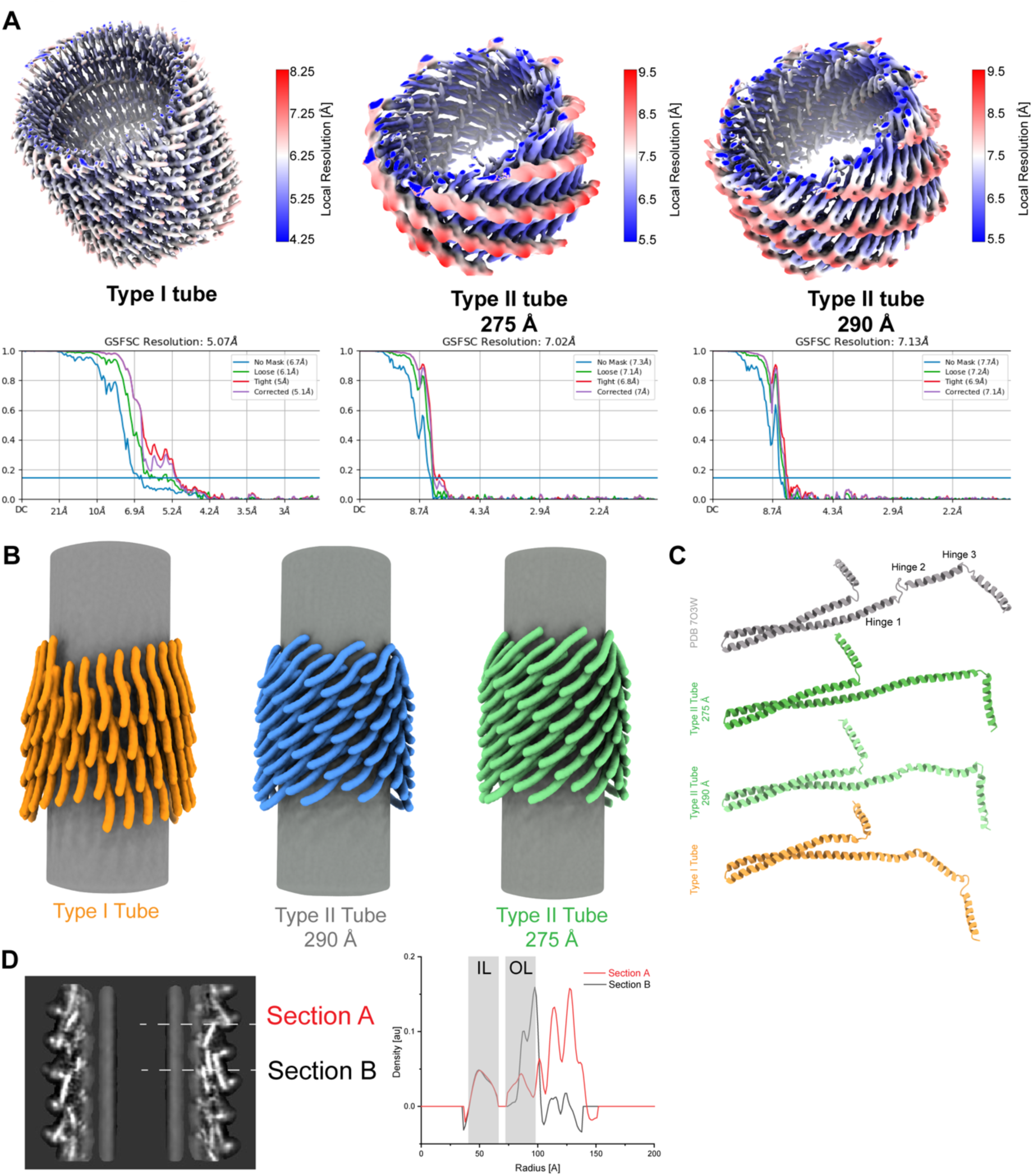
Local resolution maps and structural features of Vipp1 tubes. A: Top: local resolution maps of Vipp1 tubes (FSC=0.5). Scale bars show the local resolution range in Å. Bottom: FSC curves with global resolutions. **B:** Simplified models of the Vipp1 tubes highlighting the orientation of the monomers with respect to the membrane axis. **C:** Comparison of the monomer structures extracted from the specified Vipp1 tubes. **D:** Xy-slice of the Type I tubes with sections at the outer leaflet far from helix *α*0 (Section A) and close to helix *α*0 (Section B) and the corresponding density profiles, showing the indentation in the outer leaflet caused by helix *α*0. IL: inner leaflet, OL: outer leaflet.

**Suppl. Figure 3:**
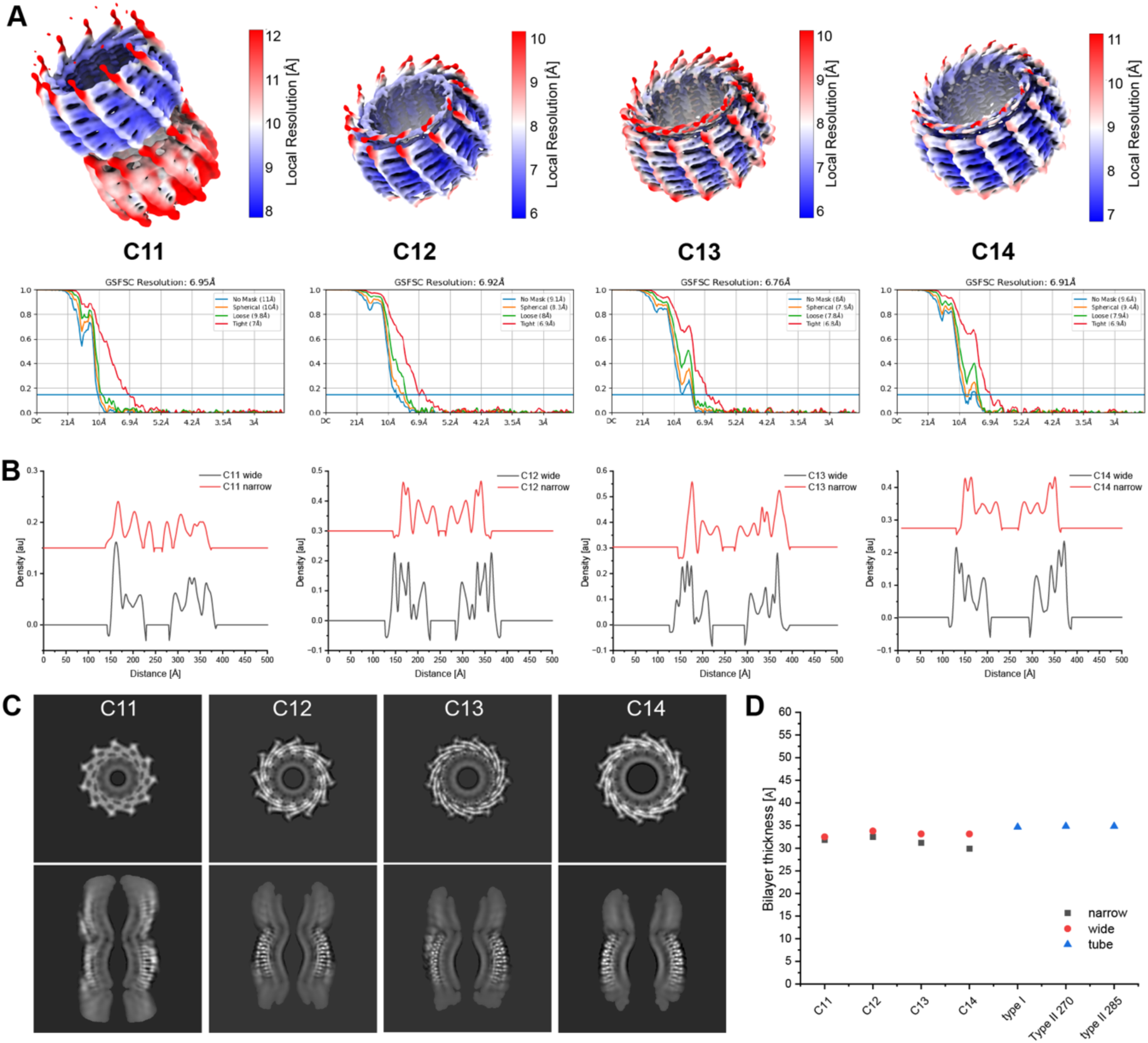
Local resolution and structural features of stacked Vipp1 rings with engulfed membrane. A: Local resolution maps of Vipp1 stacked-ring assemblies (FSC=0.5). Top: scale bars show the local resolution range in Å. Bottom: FSC curves with global resolutions. **B:** Density plots of Vipp1 rings with engulfed membrane. Red: density profile at the narrowest part of the ring; black: density profile at the widest part of the ring. **C:** Z and xy-slices of stacked C11-C14 ring models in greyscale. **D:** Scatter plot of the bilayer thickness identified in different Vipp1 assemblies.

**Suppl. Figure 4:**
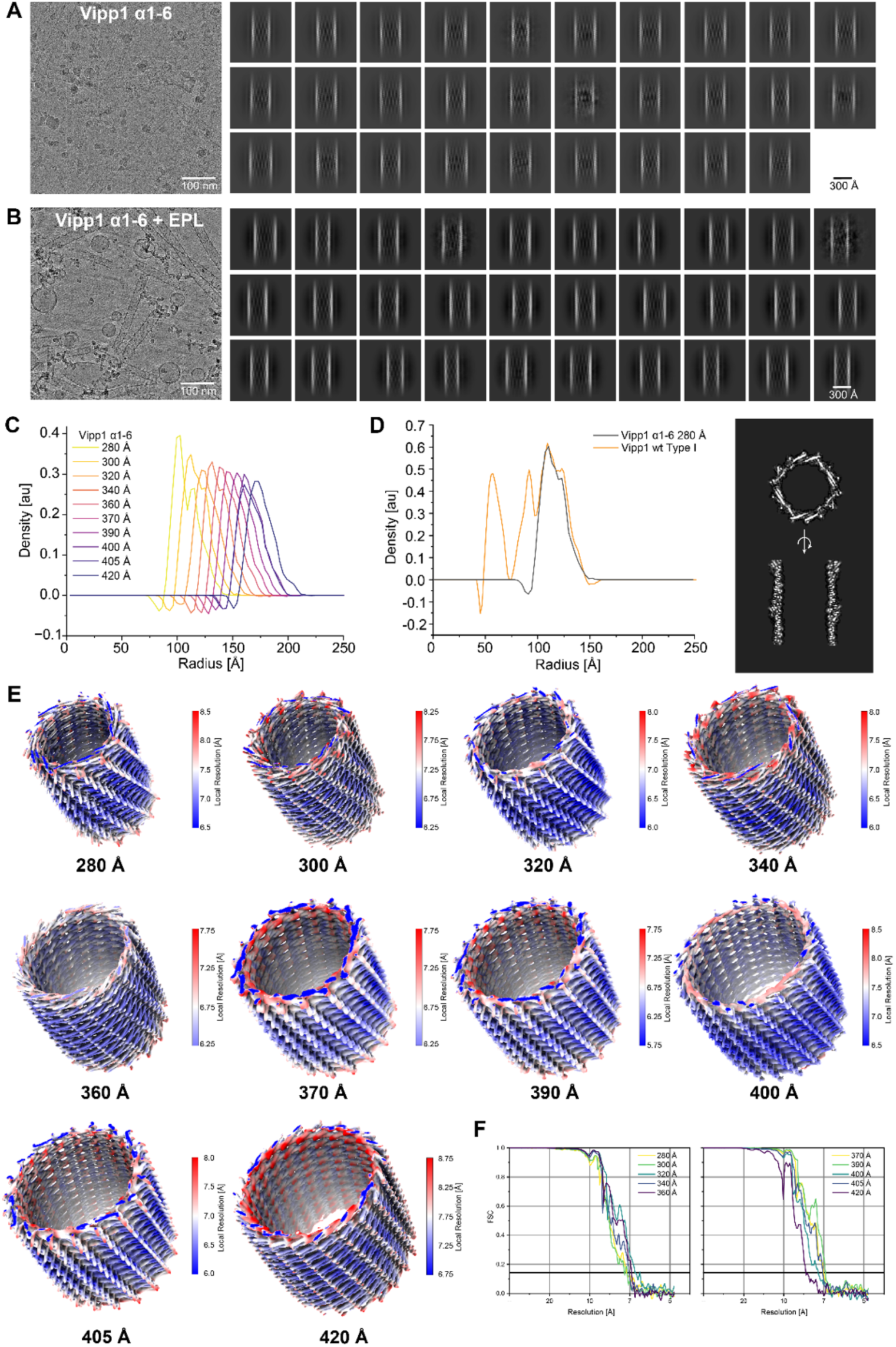
Vipp1(α1-6) helical tubes in the absence and presence of membrane. A/B: Cryo-EM micrographs and 2D class averages Vipp1(*α*1-6) with and without EPL membrane reconstitution. **C:** Radial density plots of Vipp1(*α*1–6) rods with diameters from 280 to 420 Å. **D:** Left: radial density plots of Vipp1 Type I tubes and Vipp1(*α*1–6) 300 Å rods with lipids. Right: z (top) and xy-slice (bottom) of the Vipp1(*α*1–6) 300 Å rod map in greyscale. **E:** Local resolution maps of Vipp1(*α*1-6) rods (FSC=0.5) with the corresponding diameters. Scale bars show the local resolution range in Å. **F:** FSC curves of 10 Vipp1(*α*1-6) structures with global resolutions.

**Suppl. Figure 5:**
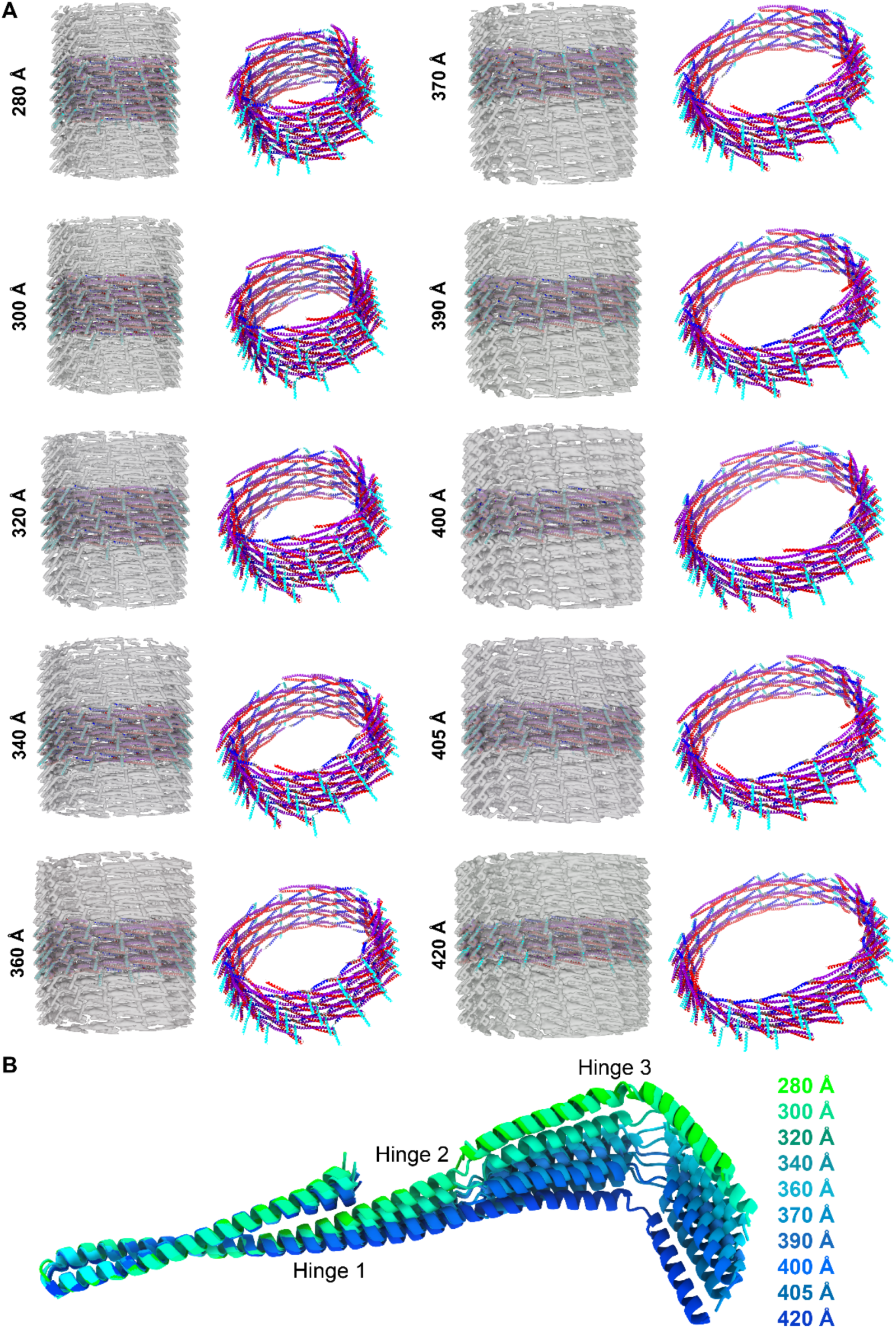
Vipp1(α1-6) helical tubes in the absence and presence of membrane. A: Cryo-EM maps with fitted models (left) and corresponding atomic models (right) of Vipp1(*α*1–6) rods in ribbon representation (*α*1 red, *α*2+3 violet, *α*4 blue, *α*5 cyan). B: Superposition of the determined Vipp1(*α*1-6) conformations with the corresponding diameters.

**Suppl. Figure 6:**
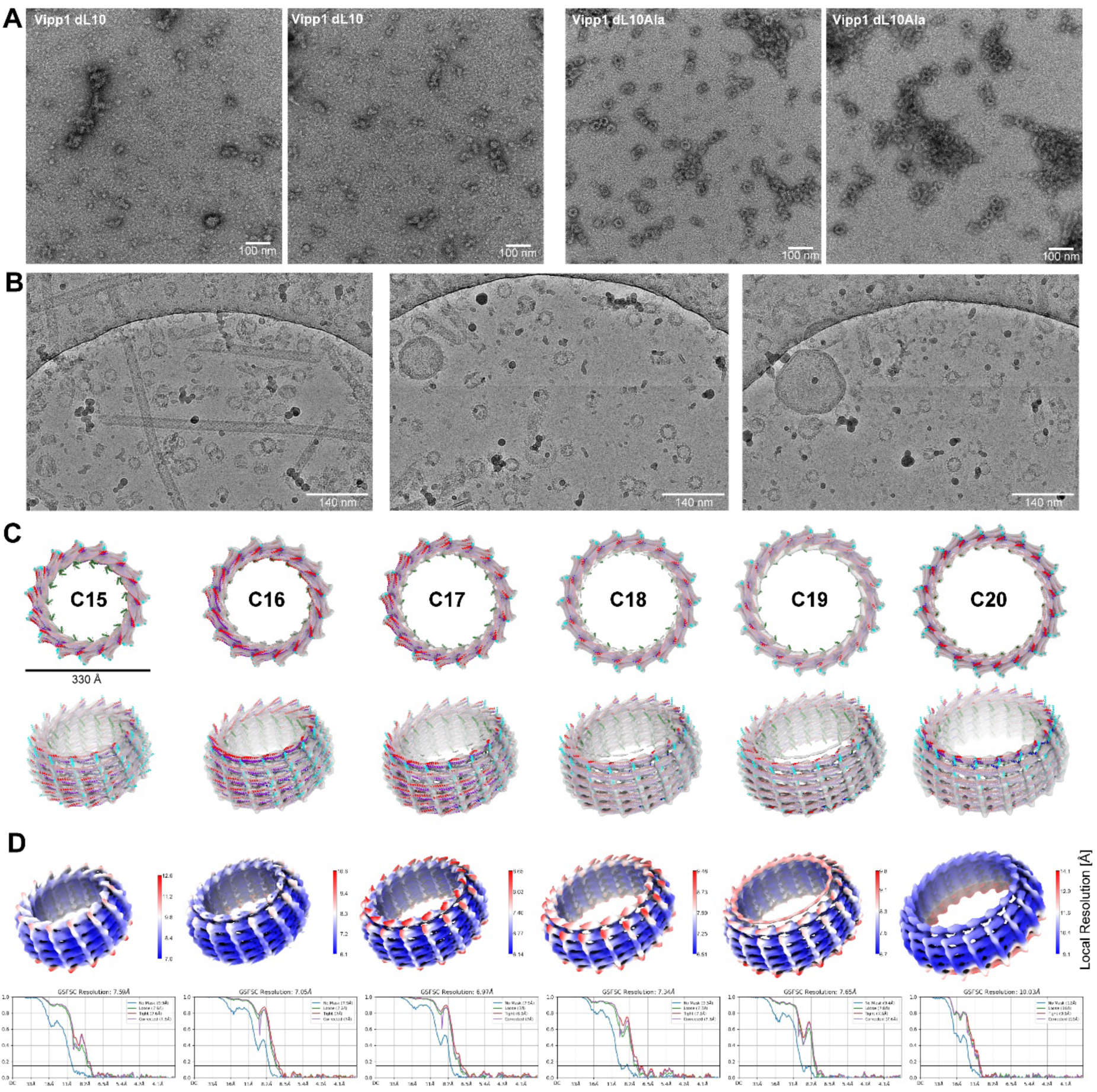
Vipp1(α1-6) helical tubes in the absence and presence of membrane. A: Negative staining EM micrographs of Vipp1dL10 and Vipp1 dL10Ala in the absence of membranes. Vipp1 dL10 forms small oligomers and larger irregular assemblies. Vipp1 dL10Ala forms small oligomers and ring complexes. **B:** Representative cryo-EM microgrpahs of the Vipp1 dL10Ala+EPL sample. **C:** Cryo-EM maps and fitted models of C15 to C20 Vipp1 dL10Ala rings. **D:** Local resolution maps of Vipp1dlA10 rings and tubes (FSC=0.5) with the corresponding diameters. Color bars indicate the mapped local resolution values in Å. FSC curves show the global resolution of the respective reconstruction at FSC=0.143.

**Suppl. Figure 7:**
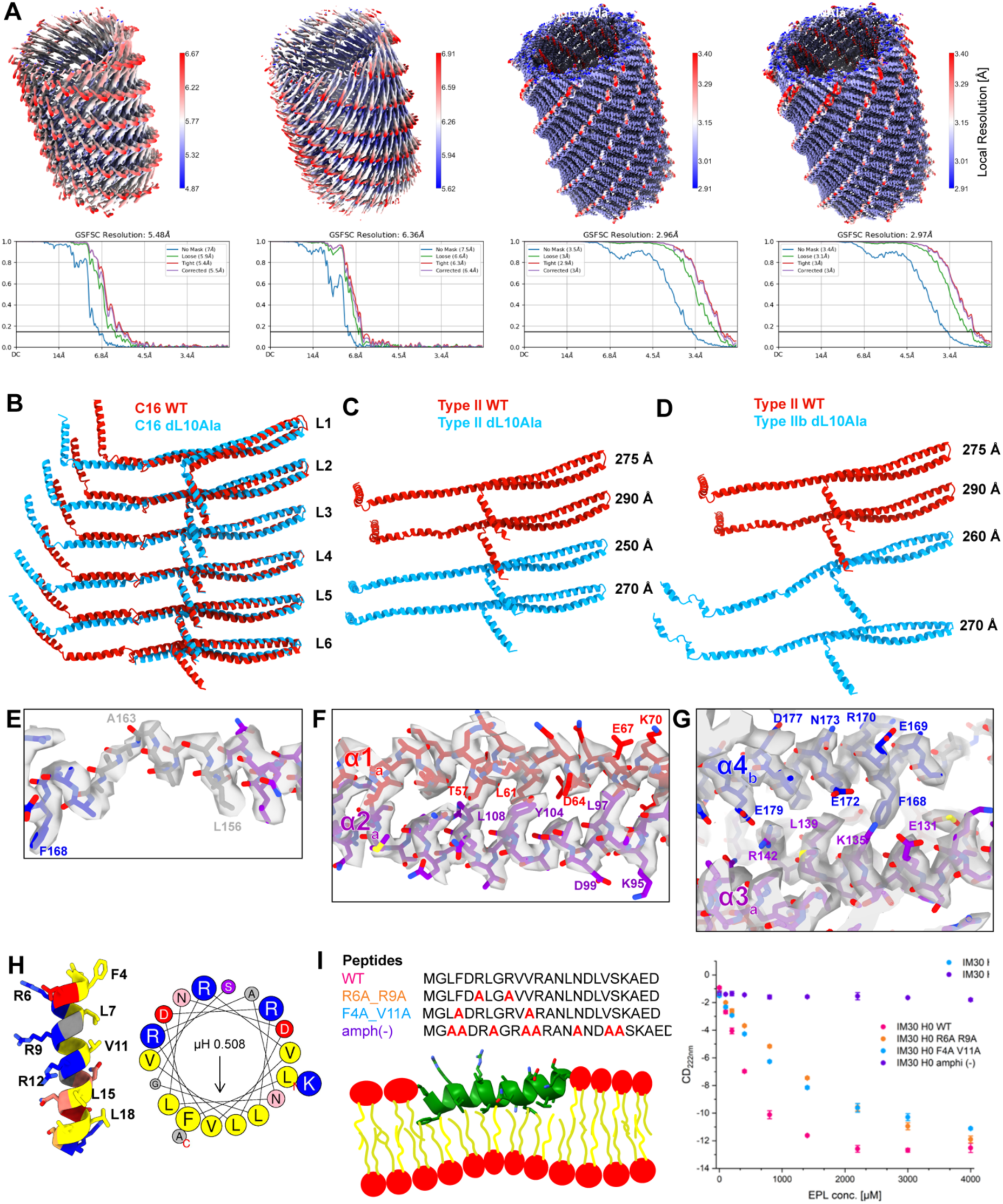
Vipp1(α1-6) helical tubes in the absence and presence of membrane. A: Local resolution maps of Vipp1dlA10 tubes (FSC=0.5) with the corresponding diameters. Scale bars show the local resolution range in Å. FSC curves show the global resolution of the respective reconstruction at FSC=0.143. **B:** Superposition of Vipp1 WT (PDB: 7O3Y) and Vipp1 dL10Ala monomers from C16 rings. **C:** Superposition of Vipp1 WT and Vipp1 dL10Ala monomers from Type II tubes. **D:** Superposition of Vipp1 WT and Vipp1 dL10Ala monomers from Type IIb tubes. **E-G:** Enlarged views of the cryo-EM map with the fitted model of Vipp1 dL10Ala Type IIb tubes. **E:** Hinge 2, **F:** Intramolecular interactions within the hairpin, **G:** Intermolecular interactions of the _168_FERM_171_ motif. **H:** Helix-wheel plot of helix *α*0 (created with HeliQuest (Gautier et al., 2008)) **I:** Membrane binding of Vipp1 helix *α*0 peptides analyzed via CD spectroscopy with the indicated mutation and putative orientation of Vipp1 helix *α*0 in the membrane, as observed in Vipp1 tubes.

**Suppl. Figure 8:**
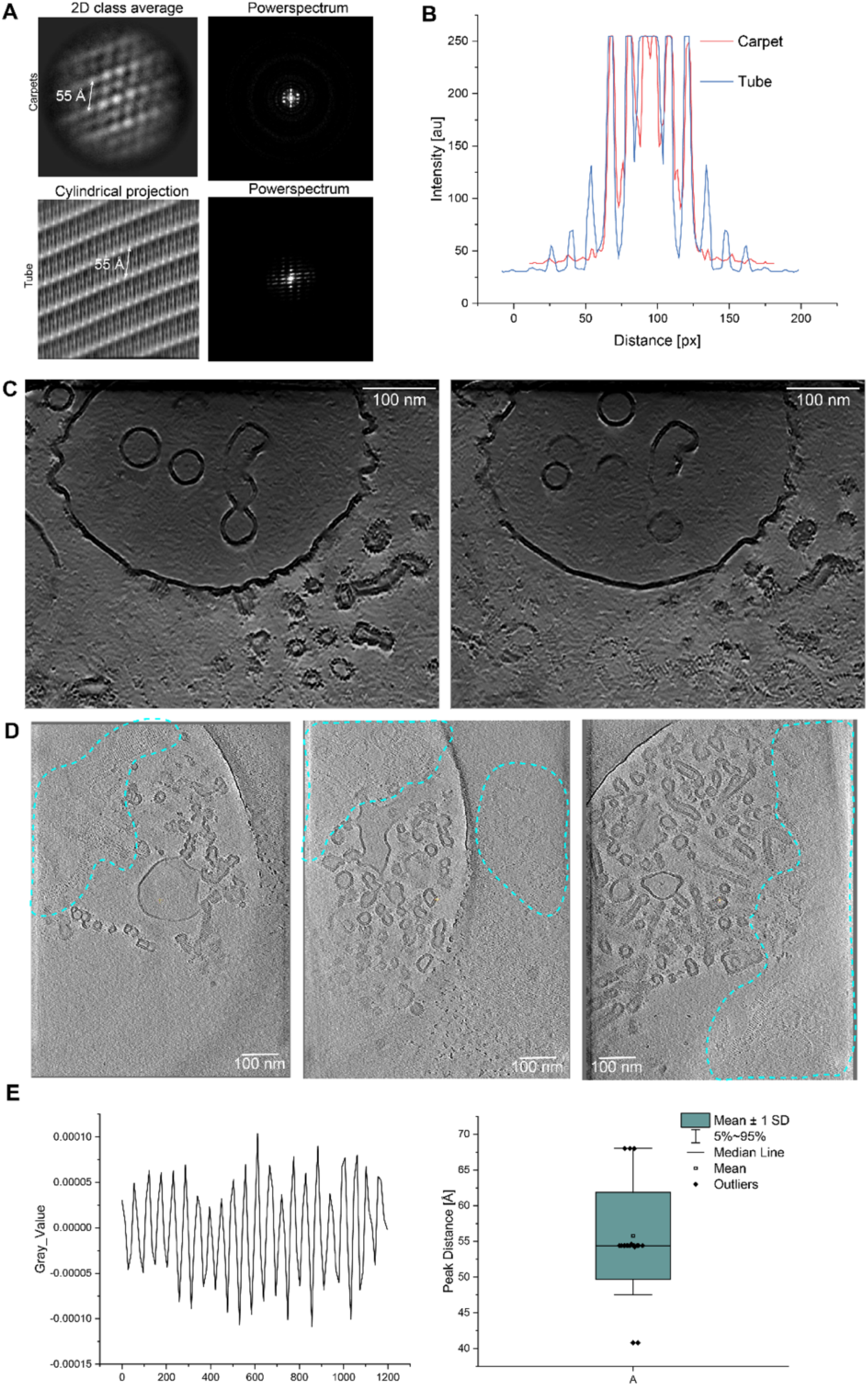
Vipp1 spirals and carpets. A: Left: 2D class averages of the Vipp1 carpets and cylindrical projection of the Type I tubes. Right: corresponding power spectra of 2D class averages of the Vipp1 carpets and Type I tubes. **B:** Overlay of the meridional density plots of the power spectra from A. **C:** Tomogram slices of Vipp1+EPL before segmentation. **D:** Tomogram slices of Vipp1+EPL. Tape-like spiral assemblies of Vipp1 are highlighted by dashed lines. **E:** Density profile of Vipp1 spirals and boxplot of the measured peak distances from the density profile, n=20.

**Suppl. Figure 9:**
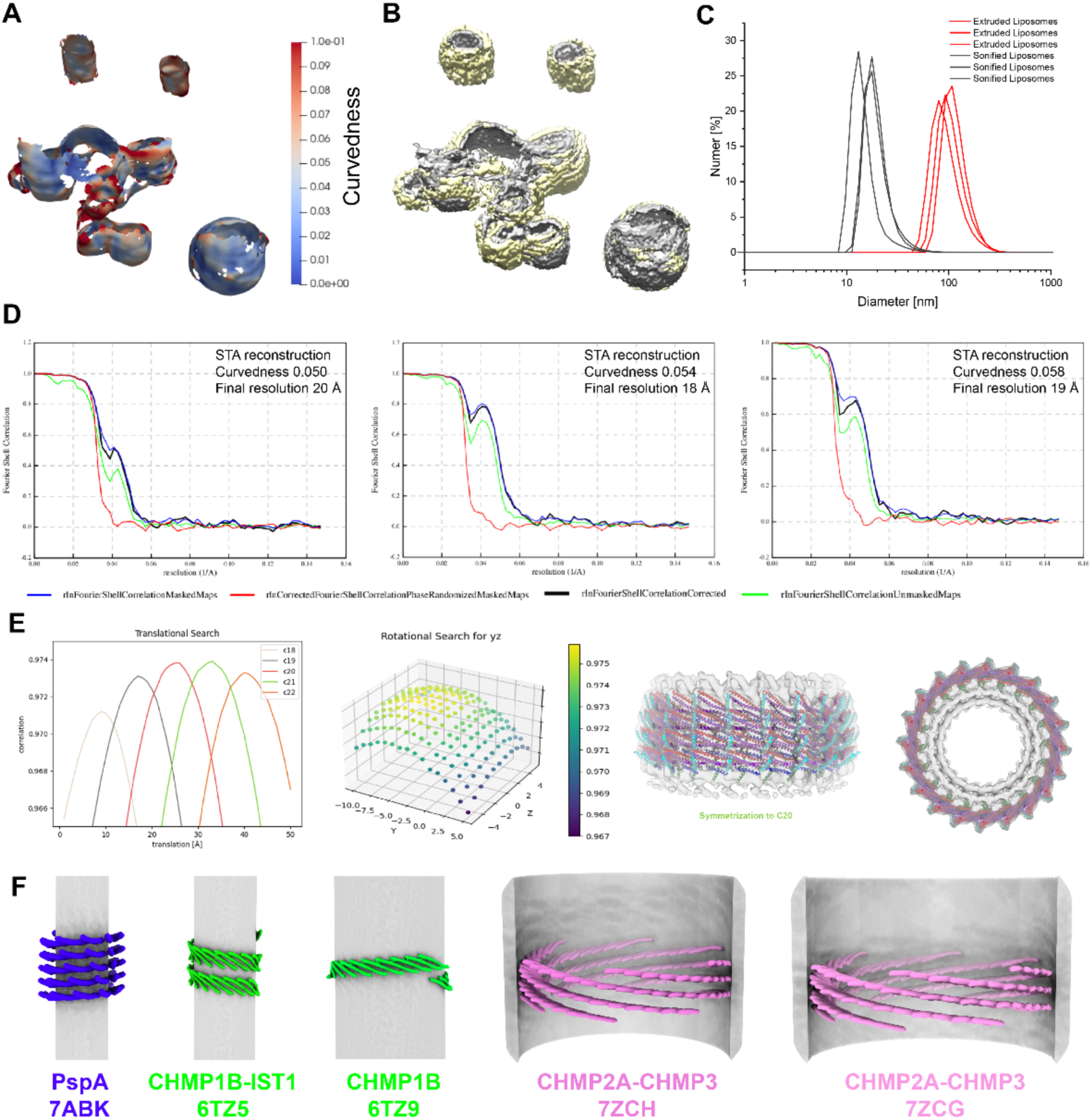
Vipp1 membrane and carpet segmentation, curvature estimation, resolution, and fitting of subtomogram averages. A: Exemplary curvedness estimates conducted with surface morphometrics. **B**: Exemplary membrane (grey) and Vipp1 carpet (yellow) segmentations. **C:** Size distribution of EPL liposomes. After preparation, the size of the liposomes was checked via dynamic light scattering. Each sample was measured three times. The results are presented as a number distribution, reflecting the fraction of particles having a given diameter. The mean size ± standard error of the number distribution is 106 ± 6 nm for the extruded liposomes (red lines) and 18 ± 2 nm (black lines). This corresponds to a diameter of 143 ± 0.5 nm for the extruded liposomes, and 56 ±0.44 nm for the sonified liposomes, based on the z-average. **D:** FSC (0.143) for subtomogram averages with curvedness 0.05, 0.054, and 0.058, respectively. **E**: Left: correlation of translational steps with different cyclical symmetries C18, C19, C20, C21, C22 (C20 yielding the highest correlation). Right: translational and rotational search (here only plotted for y and z) over different cyclical symmetries (C18, C19, C20, C21, C22) of the 0.054 curvedness structure. The C20 symmetry gave the best correlation with the subtomogram average reconstruction. Bottom row: Symmetrized subtomogram average with rigid-body fitted and symmetrized model PDB:7O3Z. **F:** Schematic views of PspA, CHMP1B-IST1, and CHMP2A-CHMP3 *α*1-3 hairpins relative to the membrane axis in stacked-ring assembly, Type II, and Type I tubes.

